# Lower Bounds on the Sample Complexity of Species Tree Estimation when Substitution Rates Vary Across Loci

**DOI:** 10.1101/2025.01.28.635331

**Authors:** Max Hill, Sebastien Roch

## Abstract

In this paper we analyze the effect of substitution rate heterogenity on the sample complexity of species tree estimation. We consider a model based on the multi-species coalescent (MSC), with the addition that gene trees exhibit random i.i.d. rates of substitution. Our first result is a lower bound on the number of loci needed to distinguish 2-leaf trees (i.e., pairwise distances) with high probability, when substitution rates satisfy a growth condition. In particular, we show that to distinguish two distances differing by length *f* with high probability, one requires *O*(*f*^−2^) loci, a significantly higher bound than the constant rate case. The second main result is a lower bound on the amount of data needed to reconstruct a 3-leaf species tree with high probability, when mutation rates are gamma distributed. In this case as well, we show that the number of gene trees must grow as *O*(*f*^−2^).

## 1 Introduction

We consider the problem of species tree estimation from genes exhibiting rate variation across the genome, that is, when mutation (e.g., substitution) rates may vary randomly between genes. In particular, we focus exclusively on rate variability between loci, as opposed to site rate heterogeneity within individual loci, or variation across lineages ancestral to a single site. We show that when data consists of error-free gene trees from the multispecies coalescent, the data requirement for accurate estimation is substantially increased compared to the case where all genes are assumed to share a common mutation rate.

The parameter we are interested in estimating is a species tree representing the evolutionary history of a set of taxa. A *species tree* 𝒮 = (*V, E, ρ, l*, ℒ) consists of a rooted leaf-labeled tree *S* = (*V, E*) with root *ρ*, branch lengths *l* =(*l*_*e*_) _*e*∈*E*_, and *n* leaves labeled bijectively by a set of species names ℒ. The edge weights *l* represent a measure of evolutionary distance and are referred to as *branch lengths*. A species tree represents a hypothesis about the evolutionary history of the species in ℒ, with edges regarded as *populations* and vertices as speciation events. In addition to the edge populations, it is also assumed that there exists a *root population*, which may be regarded as an unlabeled leaf edge of indeterminate length extending away from the root and which represents the most recent common ancestral population of all taxa in ℒ. Throughout, we shall assume that 𝒮 is *ultrametric*, meaning that all labeled leaves are equidistant from the root.

The amount of data needed to correctly estimate a species tree with high probability depends on features of the species tree. We will be interested primarily in the *minimal internal branch length f* of the species tree 𝒮, defined as

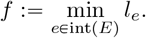

The amount of data needed to recover the species tree with high probability depends critically on this parameter due to its effect on various sources of error (including both incomplete lineage sorting and gene tree estimation error, discussed in Section 2.2). The estimation problem becomes harder as *f* becomes smaller. We are also interested in the *depth d* of the species tree, defined here as

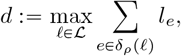

where δ_*ρ*_ (ℓ) is the set of edges in the path on 𝒮 between the leaf ℓ and the root *ρ*.

The data is assumed to take the form of *m* i.i.d. gene trees. The term *gene* is understood to refer to a non-recombining locus on the genome; that is, a gene is a specified segment of an organism’s genome which is passed down in its entirety from parent to offspring, and which contains a fixed number of DNA base pairs that themselves may mutate when this occurs. Any gene shared by the species in ℒ can be associated with a corresponding *gene tree* 𝒯, a rooted edge-weighted tree with leaves labeled bijectively by ℒ. A gene tree is regarded as representing the genealogical history of a single gene, which in general may differ from the population-level evolutionary history represented by 𝒮. The leaves of 𝒯 are regarded as members of a homologous gene family, with each member having been sampled from a species in ℒ. Unlike the edges of the species tree, which represent populations, the edges of a gene tree are regarded as *ancestral lineages*, that is, lines of descent for particular sampled individuals, and the vertices of 𝒯 correspond to most recent common ancestors.

The nexus question of this paper pertains to how many gene trees are needed—as a function of *f* and *d*—in order to have a high probability of correctly estimating an unknown species tree parameter as (*f, d*) → (0, ∞). Our main result is an information-theoretic lower bound which says that, independently of *d*, the number of sampled gene trees *m* must grow at least as quickly as *f* ^−2^ when genes exhibit gamma-distributed random mutation rates. This is a substantially greater data requirement than is the case when constant mutation rates are assumed, as in that case a data requirement of *m* = *O*(*f* ^−1^) gene trees is known to be sufficient (see Section 2.2). The lower bound we prove is tight, since a corresponding upper bound of *m* = *O*(*f* ^−2^)is known to be achievable (see Fig. 1).

**Figure 1:**
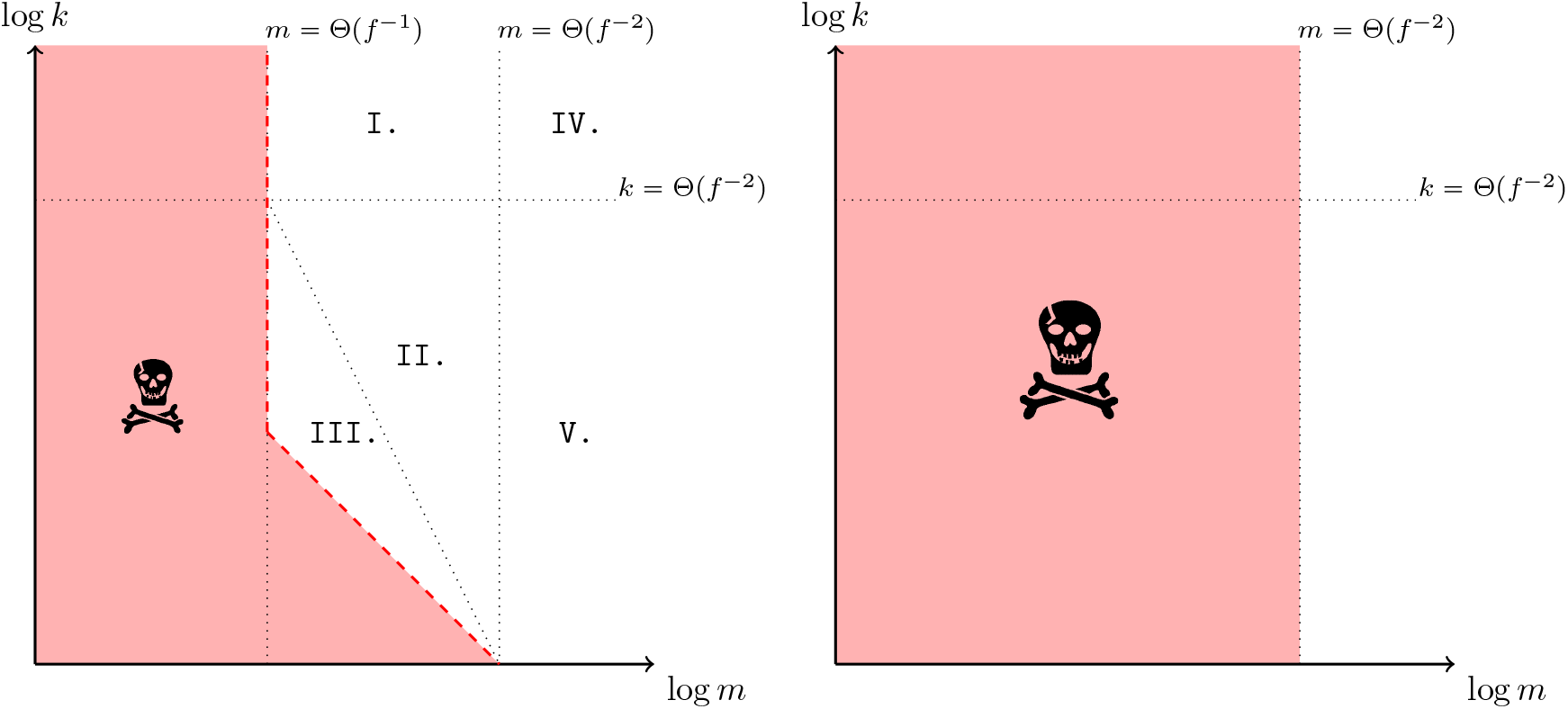
*Left:* A cartoon log-log plot summarizing the known sample complexity results for species tree estimation from sequence data under the assumption that mutation rates do not vary across genes, as *f →* 0. The red dashed line corresponds to Eq. (4), and the shaded region marked with 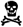 is the impossibility zone, where consistent inference is not possible by any method. Upper bounds are achieved by GLASS (in region `I`), `QuEST` (in region `II`), `STEAC` and topological inference methods like `ASTRAL` and `SEQUINED` (in region `IV`), and `METAL` and `fmulti-sites` [1] (in region `V`). Methods achieving consistent inference in one region also achieve it in all regions where more data is available (viz. above or to the right). The line separating regions `II` and `III` corresponds to Eq. (5); in region `III`. In region `III`, it is known that distance-based methods cannot achieve consistent inference [2] and it is suspected that inference is not possible by any means. *Right:* The sample complexity when mutation rates are allowed to vary independently between genes according to a Gamma distribution (possibly with an atom at zero). The dramatic expansion of the impossibility zone and subsequent collapse of the information-theoretic tradeoff between *m* and *k* described in Section 2.2.2 is the main result of this paper.

This paper is structured as follows. In Section 2, we review related work in models of rate variation and sample complexity in phylogenetics, introduce our model of gene evolution, and summarize the general approach that we will take to prove our main results. In Section 3 we prove the first main result, which considers the special case of distinguishing edges when mutation rates are gamma-distributed, along with an instructive comparison to the case of nonrandom mutation rates. In Section 3.4, we describe a complication which arises when attempting to generalize the first main result to more general probability distributions. In Section 4, we state and prove our second main result, a lower bound on the amount of data necessary to achieve high probability of correctly distinguishing a binary 3-leaf species tree from a star tree.

## 2 Background

### 2.1 Rate variation models

The standard model of rate variation across the genome is a mixture model known as the *rates-across-sites* (RAS) model (see e.g., [3, 4]). In the RAS model, the underlying tree topology is assumed to be the same for all sites, but the mutation rate for each site is chosen randomly in an i.i.d. manner from a distribution, resulting in branch length variation between sites. The most widely-used formulation is the *general time-reversible model with gamma-distributed rates and invariable sites*, abbreviated GTR + Γ + I. Under this model, each site is either invariant or undergoes a mutational process governed by a continuous-time Markov process with rate matrix *rQ*, where *Q* is a general time-reversible instantaneous rate matrix, and *r* is an independent gamma-distributed random variable with rate *α* > 0 and shape *β* > 0. Both *α* and *β* may be regarded as unknown parameters, though typically *β* is chosen to be 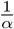 so that *r* has mean 1. In another widely-used formulation, *r* is assumed to have a discrete gamma distribution [5, 6].

When *α* is unknown, the question of identifying model parameters (which include *Q, α*, the stationary distribution, and all features of the species phylogeny) is somewhat subtle, as pairwise sequence comparisons are not sufficient to recover the model parameters [7]. Nonetheless, it has been shown that all parameters are generically identifiable from sequence data under both the GTR + Γ model [4], as well as the GTR + Γ + I model under reasonable assumptions about the species tree branch lengths and rate matrix *Q* [8]. Moreover, the species tree topology is generically identifiable under GTR + Γ + I even when the assumption that all sites share a common tree topology is relaxed, as in the coalescent-based model of [9]. More general RAS models were considered by [10] in the setting of the *large-tree limit* (i.e., as the number of leaves *n* tends to infinity), who showed that for sufficiently large trees, the topology of the species tree is both identifiable from the distribution of sites and can be recovered with high probability from sequence data of polynomial length in *n*.

The model of rate variation considered in this paper will differ from RAS models in an important way: rather than requiring that all gene trees share a common topology, it will instead be assumed that gene trees are generated according to the *multispecies coalescent* (MSC) [11], with mutation rates randomly picked for each gene. To our knowledge, this is the first paper to establish theoretical bounds on the sample complexity of phylogenetic inference for coalescent-based models involving rate variation between genes, and as we will show, the impact of rate variation on the sample complexity is significant. To frame our main results, it will be helpful to review what is known about the sample complexity of coalescent-based models.

### 2.2 Sample complexity in phylogenetics

Broadly speaking, the term *sample complexity* refers to the amount of data needed to achieve high probability of correct estimation. For coalescent-based models, this has been studied from two perspectives: (1) using gene trees as data, and (2) using sequences as data. The common assumption in case (1) is that gene trees are drawn according to the MSC in an i.i.d. manner and that one has full information about each gene tree. For case (2) on the other hand, a 2-step model is typically employed. First, *m* gene trees are drawn independently according to the MSC. Then, *m* sequences of length *k* are generated independently by a Markov model of site substitution (e.g., the Jukes-Cantor model) using the *m* respective gene trees as input. Unlike in the first case, the gene trees are assumed to be unknown; only the sequences are observed.

In both cases, the aim is to estimate the topology of an unknown species tree parameter, and the key question pertains to how the data requirement for consistent inference grows as a function of certain key model parameters. This question has two aspects: *lower bounds*, which pertain to how much data is *necessary* for any inference method to succeed with high probability, and *upper bounds*, which pertain to how much data is *sufficient* for a particular inference method to succeed with high probability. Lower bounds provide both a measure of the overall difficulty of the estimation problem, while upper bounds provide a criterion by which to compare the performance of difference inference methods. The sample complexity of species tree estimation is analyzed in the asymptotic regime in which one or more structural parameters (e.g., tree depth, minimal branch length, number of leaves) are approaching some limits.

In case (1), the data requirement can be expressed by a single quantity, *m*, the number of gene trees. In case (2) by contrast, the total data is the product *mk*, where *k* is the length of each gene sequence *k*. Of course, for real-world data the total sequence length *mk* is constrained, which results in an information-theoretic tradeoff between *m* and *k*. Much of the work in this area has been aimed at better understanding how inference is affected by this tradeoff.

In particular, what makes this tradeoff nontrivial is that the quantities *m* and *k* do not play equivalent roles. Increasing one is not equivalent to increasing the other, because they are related to very different sources of error. On one hand, when *k* is large, the accuracy of each estimated gene tree should increase, so that one may need fewer gene trees to estimate the species tree [2]. On the other hand, it is possible to have too few gene trees, irrespective of how large *k* may be. This is because observing a sufficient number of genes is necessary to mitigate the risk of error arising from *incomplete lineage sorting* (*ILS*), a phenomenon in which gene lineages fail to coalesce within a population, which can result in topological discordance between gene trees and species tree as well as a failure to detect short internal branches of the species tree [2, 12, 13, 14].

To accompany this review, a graphical summary of the known sample complexity results for species tree estimation is shown in Fig. 1.

#### 2.2.1 Sample complexity for gene tree data

The first fundamental data requirement for consistent species tree estimation from gene tree data is that as *f* → 0 it is necessary for

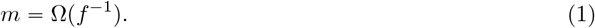

This lower bound (see [2]and Appendix A), stems from the fact that achieving high probability of correct inference is only possible if *at least one gene tree does not exhibit ILS on the shortest internal branch of the species tree*, an event which we will denote by. 𝒜 Only if 𝒜 occurs does the data provides any evidence for the existence of the short branch. The `GLASS` algorithm, a distance-based method which also uses minimum coalescent times from the *m* gene trees, but which can be used to estimate trees with any number of leaves, has been shown to return the correct tree topology with high probability provided that *m*= *O*(*f* ^−1^) [15]. Eq. (1) implies that this is the best achievable bound in the limit as *f*→ 0, up to multiplication by a constant.

The `GLASS` method is somewhat atypical in that it achieves the lower bound of *m*= *O*(*f* ^−1^); the upper bound of *m*= *O*(*f* ^−2^)is more common, especially among methods used in practice. Another algorithm which has been well-studied is `STEAC` (Species Tree Estimation using Average Coalesecent times [16]), which utilizes *average* rather than *minimum* coalescent times across gene trees. Using gene trees as data, `STEAC` achieves consistent inference with *m*= *O*(*f* ^−1^)[17]. This is a much higher data requirement when *f* is small. Another important class of algorithms are *topological inference methods*, which make use of only the gene tree topologies, not the branch lengths. Many of these are *quartet-based methods*, which take the approach of estimating unrooted four-taxon trees and then combining these estimates in some manner to reconstruct the full species tree. For such methods, the sample complexity upper bound of *m*= *O*(*f* ^−2^)is known to be achievable. For example, `SEQUINED`, a naive quartet method based on topological distances^1^ between taxa; `SEQUINED` succeeds with high probability if *m*= *O*(*f* ^−2^)[18]. Another example is a suite of methods known as `ASTRAL` [19], which estimate the species tree from a collection of gene trees by scoring trees based on how many unrooted topological quartets they share with the gene trees. In [20], it was shown that when *f* is small, `ASTRAL` achieves high probability of correctly estimating the species tree topology if and only if that the number of samples is at least *m*= Ω (*f* ^−2^). A similar approach to the upper bound of [20] was used in [21] to show that this upper bound also holds for gene trees generated under the DLCoal model of gene duplication and loss introduced in [22]. A different asymptotic regime is considered in [23], which focuses on the probability of incorrect tree estimation as *m*→ ∞ for a fixed species tree *S*; in that paper it is shown that for a general model of evolution which includes the MSC, the error probability of quartet-based methods decays exponentially as *m m*→ ∞, and upper bounds are numerically obtained which improve on those in [20] when *m* is large.

#### 2.2.2 Sample complexity for sequence data

So far we have discussed sample complexity bounds under the assumption that one has access to error-free gene trees. In practice however, gene trees cannot be observed directly but rather must be inferred from DNA sequence data. Moreover, using sequence data to estimate trees introduces a second source of error apart from ILS—namely, the stochastic nature of the substitution process.

Any method for estimating trees from sequence data under models involving a Markov site substitution process are subject to a *sequence length requirement*, which corresponds to the number of independent sites the method needs to accurately estimate the unrooted tree topology with high probability [24]. For sequence data under a site substitution model alone, not including ILS as a source of error, a lower bound on the sequence length requirement grows proportionally to the inverse square of the minimal branch length [25]. Thus, to estimate a tree not subject to ILS (e.g., a gene tree) correctly with high probability from a sequence of length *k*, it is necessary that

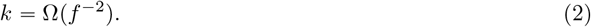

Correspondingly, it is also known that *m*= *O*(*f* ^−2^)is sufficient to achieve high probability of successful estimation [26, 27]. Hence, inference methods which assume that the data take the form of *m* error-free gene trees from the MSC can be extended to sequence data using estimated gene trees. In particular, if *k* Ω (*f* ^−2^)then we expect such methods to succeed with high probability provided that their data requirement for *m* is met (e.g., that *m* scales as *f* ^−1^for `GLASS` or as *f* ^−2^for `STEAC` and topological inference methods).

The aforementioned lower bound on the sequence length requirement also holds as a lower bound for coalescent-based models, since the inclusion of ILS as an additional confounding source of error can only make the estimation problem harder. Thus, in the case of 2-step models which combine the MSC with a Markov substitution process to produce data taking the form of *m* independent gene sequences each of length *k*, the sequence length requirement translates to the lower bound that the total sequence length *mk* must satisfy

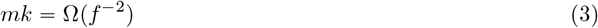

as *f* → 0 [12].

In addition to the above sequence length requirement, the number of genes *m* must also satisfy *m* =Ω (*f* ^−2^)independently of *k*. This is because ILS represents a source of error which cannot be overcome solely by increasing the lengths of individual gene sequences. Consequently the lower bound in Eq. (1) holds for sequences of any length [12]. Taken together, Eqs. (1) and (3) imply that in order to correctly estimate a species tree with high probability from sequences of length *k* ⩾1, the number of genes *m* must satisfy

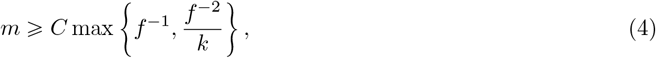

for some constant *C >* 0 not depending on *f* or *k* [12].

The form of Eq. (4) suggests that the data requirement for *m* may depend markedly on the growth rate of *k*, and indeed this is the case: there is a spectrum of very different regimes in which species tree estimation is possible, ranging from the small *k* =*O* (1) to the very large *k* =Ω (*f* ^−2^). In the latter case where *k* is very large, the upper bound *m*= *O*(*f* ^−1^)is notably achieved by `GLASS`, as discussed in Section 2.2.1.

On the other hand, when *k* =*O* (1), it follows from Eq. (4) that *m*= Ω (*f* ^−2^)necessary for consistent inference. Moreover, the lower bound in Eq. (4) is achievable when *k* =*O* (1), by a `STEAC`-like algorithm known as `METAL` (Metric algorithm for Estimation of Trees based on Aggregation of Loci [12, 28]); in the `METAL` algorithm, each taxon’s gene sequences are concatenated together into a sequence of length *mk*, then normalized Hamming distances are computed between these concatenated sequences for each pair of taxa, and a clustering algorithm is applied. For any *k* ⩾ 1, `METAL` succeeds with high probability as long as *m* scales like Ω(*f* ^−2^) [12, 28].

There is a subtle information-theoretic tradeoff between *m* and *k* in the spectrum between these two extremes. This tradeoff has been carefully studied for *distance based methods*, i.e., methods relying only on the number of pairwise differences between gene sequences for each pair of taxa (e.g., `METAL`) [2]. The sample complexity of distance-based methods in the intermediate zone between the two extremes of *k* =*O* (1) and *k*= Ω (*f* ^−2^)was explored in [2, 29], where it was shown that for to achieve high probability of correct inference as *f* → 0 using distance-based methods, it is necessary and sufficient that

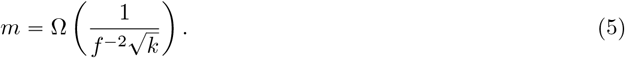

That this many samples is sufficient for consistent estimation was demonstrated via the introduction of a novel distance-based method known as `QuEST`, which utilizes a quantile comparison of distance estimates [2, 29]. In Fig. 1, Eq. (5) corresponds to the dotted line separating regions `II`. and `III`.

### 2.3 Our Model

#### 2.3.1 Mutation rates and measurement of evolutionary distance

There is an important difference between the branch lengths of the gene trees and the branch lengths of the species tree: they do not use the same units. For the model of gene evolution which we will adopt in this study, the mutation rate will be taken to be a random quantity which varies from gene to gene. The appropriate unit of measurement for gene tree branch lengths in this context is *expected number of mutations per site*. By contrast, since the branch lengths of the species tree are *model parameters* (which ipso facto must be chosen at the outset), it is not possible to specify them in terms of the expected number of mutations per site, as this unit of measurement necessarily depends on the mutation rate, which is random. Instead, we assume that the branch lengths of the species tree are measured at the outset in *coalescent units*, where

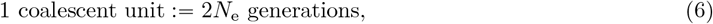

and *N*_e_ is the effective population size.

The *scaled mutation rate* is defined as

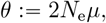

where *µ* > 0 is the rate of mutation per site per generation. Use of the combined parameter *θ* is generally preferred since *N*_e_ and *µ* are not separately identifiable from DNA sequence data alone [30, 31].

Given a gene which undergoes spontaneous neutral mutation at a rate of *µ* mutations per site per generation, it follows by Eq. (6) that the expected number of mutations per coalescent unit is *θ*. Hence we can measure evolutionary distance (i.e., branch lengths of a species or gene tree) alternatively by *coalescent units* or by *expected number of mutations per site*, converting between the two units of distance by means of a multiplicative factor:

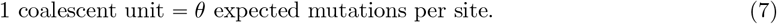

In this paper we assume that the scaled mutation rate *θ* is a random variable chosen drawn independently for each gene according to a pre-specified probability distribution *µ*_*θ*_, but that mutation rates do not otherwise vary in time or across populations. A commonly used distribution for modeling mutation rates is the gamma distribution, or a mixture of a gamma distribution with a Dirac point mass at zero (i.e. an “atom” at zero) [4, 32, 33], which is what we shall consider in this paper. To be precise, a random variable *G* is said to be *gamma-distributed*, denoted *G* ∼ Γ(*α, β*), if the distribution *µ*_*G*_ of *G* has probability density function 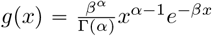 with support (0, ∞) for some *α, β* > 0. And we say that *θ* is *gamma-distributed with an atom at zero* if there exists a constant λ ∈ (0, 1) such that

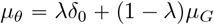

where δ_0_ is the Dirac measure at 0 on ℝ, or in other words, if

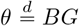

where Λ is an independent random variable with

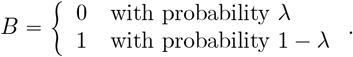

The justification for the widespread use of a gamma distribution is rather thin—resting mostly on mathematical tractability—however it is understood that the consideration of a point mass at zero is important to model genes which are invariant, for example genes which are sufficiently conserved that they do not change at all over long periods of time [32].

#### 2.3.2 Multispecies Coalescent (MSC)

To model gene evolution, we will use the *multi-species coalescent* (MSC), due to [11], which describes a probability distribution of a gene tree *T* as a function of a species tree parameter [31], and which may be regarded as an algorithm for generating gene tree samples from a species tree [30].

The MSC can be described constructively as follows: one starts with some number of gene copies at the leaves of 𝒮 (e.g., one gene copy in each species of ℒ_*S*_), and then traces each of these copies upwards within the edges of 𝒮, forming lineages going backwards in time toward the root of 𝒮. Any pair of lineages located in the same edge of 𝒮 are regarded as being in the same population, and pairs of lineages in the same population *coalesce*, or join together into a single lineage, independently at rate 1 /*θ*. The process continues until all lineages have coalesced and only a single lineage remains, an event which occurs with probability one in the root population of 𝒮. The output is a gene tree representing the genealogy of the gene copies initially sampled from the leaves of 𝒮. A fuller descriptions of this model can be found in [11, 31, 14, 30].

For our present purposes we will need a precise definition of the density function of a gene tree 𝒯 under the multispecies coalescent. As shown in [11, 31], this can be obtained by factoring the density of the full tree into densities across the populations of *S* in the following way.

Let *V* be the set of vertices in 𝒮, and for each *v* ∈ *V* denote by *τ*_*v*_ the age of vertex *v* (in coalescent units). For each *v* ∈ *V* \ {*ρ*} let 𝒳_*v*_ be the population in 𝒮 corresponding to the edge (*v*, pa(*v*)) in 𝒮, where pa(*v*) denotes the parent vertex of *v* relative to the root. If 𝒯 has *m*_*v*_ lineages entering population 𝒳_*v*_ at time *τ*_*v*_ and *n*_*v*_ lineages exiting 𝒳_*v*_ at time *τ*_pa(*v*)_, let 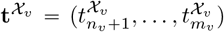, be the vector of *m*_*v*_ − *n*_*v*_ coalescent waiting times, where 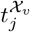 is the waiting time in population 𝒳_*v*_ until the number of lineages drops from *j* to *j* − 1 (which occurs when a pair of lineages coalesces), with 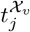 measured in expected number of mutations per site. Let 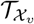 denote the portion of *𝒯* restricted to population 𝒳_*v*_. Given a scaled mutation rate of *x* > 0, the probability density of 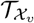 is

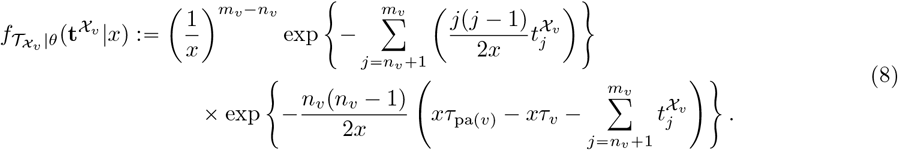

For the root population *𝒳*_*ρ*_, which starts at time *τ*_*ρ*_ and extend indefinitely into the past, we must have *n*_*v*_ − 1, and the density of 𝒯 in 𝒳_*ρ*_ is given by the same formula in Eq. (8), with the second exponential term having vanished. The probability density function for the whole gene tree with coalescent waiting times 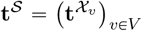 is then given by the product

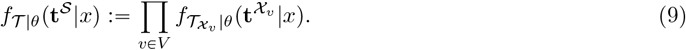

In the case that a gene tree *𝒯* has distribution given by Eq. (9) with scaled mutation rate *x* > 0, we write

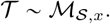

In words, *ℳ*_*S,x*_ is the distribution of a gene tree generated according to the MSC on *𝒮* with a given fixed scaled mutation rate *x*> 0. For a random *θ*> 0 with probability distribution *µ*_*θ*_, the density function *f*_*𝒯*_ of the gene tree is

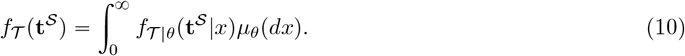

We denote the distribution given by this density function as 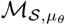.

This formula will be sufficient for the case where *θ* is Gamma distributed. However, it is not sufficent for the case where *θ* has an atom at zero, since *f*_*𝒯* |*θ*_(· |0) is undefined. The simplest way to extend the model to allow for *θ* to have an atom at zero is through the use of a scaling factor. A gene exhibiting zero mutation rates is assumed to be noninformative; let *T*_0_ denote the gene tree with zero branch lengths and star topology. Given a gene tree 𝒯 and a nonnegative number *x*, define *x* · *𝒯* to be the tree obtained by multiplying all branch lengths of *𝒯* by *x* if *x* > 0, and *T*_0_ if *x* = 0.

Let *µ*_*θ*_ be a probability distribution (not necessarily continuous) supported on [0, ∞), and consider the following procedure:

1. Draw a tree 𝒯 according to ℳ_*S*,1_. (Recall that _*S*,1_ is the special case of the MSC where the base rate of coalescence for each pair of lineages in a population is 1).
2. Independently, draw *θ* according to *µ*_*θ*_.
3. Return the tree *θ* · 𝒯.

The next lemma shows that when *µ*_*θ*_ has a density function *g* such that *g*(*x*) = 0 for all *x ⪕*0, the above procedure generates a gene tree with distribution 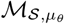 (i.e., it agrees with the gene tree distribution under the MSC as defined in the previous section).

##### Lemma 1.

*Let* 𝒯 ∼ *ℳ*_*S*,1_, *and let θ be a random variable with distribution µ*_*θ*_ *having probability density function g such that g*(*x*) = 0 *whenever x ⪕*0. *Then θ* · 𝒯 *has distribution* 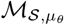.

A proof is provided in Appendix B.

In contrast to the model description given in Section 2.3.2, the scaling procedure described here makes use of only one assumption about the random variable *θ*—namely, that it is nonnegative. Therefore, in light of Lemma 1, we may *extend* our previous definition of 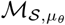 using the scaling procedure to admit any probability distribution *µ*_*θ*_ supported on [0, ∞). In particular, we define 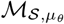 to be the distribution of the random gene tree *θ* · 𝒯 where 𝒯 is drawn according to ℳ_*S*,1_ and *θ* is drawn independently according to *µ*_*θ*_. We will require this definition when considering mutation rates which have an atom at zero.

### 2.4 Key ingredients for proving sample complexity bounds

In this section we describe the general tools relating to total variation and Hellinger distance as they are applied to a hypothesis testing problem like that considered in this paper. The proofs of our main results will follow the approach of Theorem 1 in [2], which used a bound on the Hellinger distance to bound the total variation distance, which in turns yields a lower bound on the number of samples needed for consistent species tree estimation. We explain this approach in general terms here.

#### 2.4.1 Total variation distance

For probabiliy measures *µ* and *ν* defined on the same measurable space, the *total variation distance* is defined as

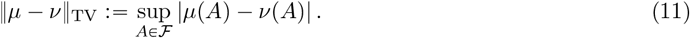

The total variation distance provides a way to measure the difficulty of a hypothesis testing problem, as we now describe. Let *P* and *Q* be probability measures defined on the measurable space (Ω, ℱ), and let *P* ^⊗*m*^ and *Q*^⊗*m*^ denote distributions of *m* independent samples from *P* and *Q* respectively. Consider the following hypothesis testing problem:

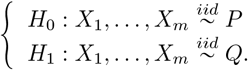

The goal is to distinguish *P* and *Q* on the basis of *m* independent samples drawn from one of the distributions.

Any statistical test for doing so can be identified with its *rejection region*, a measurable subset set 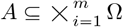 defined as the set of sample points for which *H*_0_ is rejected [34]. Given any test *A*, the maximum of its probabilities of type-I and type-II errors is

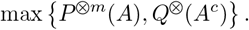

Taking the infimum over all tests,

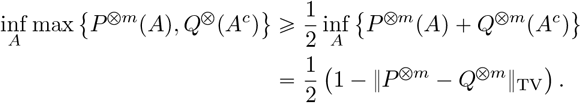

Hence if ‖*P* ^⊗*m*^ − *Q*^⊗*m*^‖_TV_ ⩽*ϵ*, then any statistical test to identify tree parameter from *m* samples drawn from one of the distributions *P* or *Q* cannot have type-I and type-II error probabilities both less than 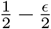. (For a detailed reference, see Theorem 2.2 in [35].) Therefore, to establish a lower bound on *m* needed to distinguish *H*_0_ from *H*_1_, it is enough to bound

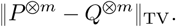

#### 2.4.2 Hellinger distance

Given two probability measures *P* and *Q* defined on (Ω, ℱ), the *Hellinger distance*, denoted *H*(*P, Q*) is defined the *L*^2^-distance between the square roots of their densities; that is,

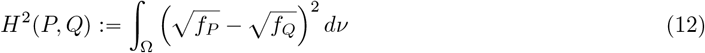

where *ν* is any measure such that the densities *f*_*P*_ and *f*_*Q*_ are absolutely continuous with respect to *ν*.

The Hellinger distance satisfies two important properties which we will make use of. The first is that

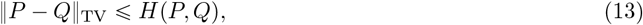

which follows from the Cauchy-Schwarz inequality (see, e.g., [36]). The second key property, stated in the following lemma, generalizes the first. It relates the total variation distance between the product measures *P* ^⨂*m*^ and *Q*^⨂*m*^ (i.e., the distribution of *m* independent samples from *P* and *Q* respectively) to the squared Hellinger distance of *P* and *Q*.

##### Lemma 2

(Hellinger bound for product measures). *Let P, Q be two probability distributions defined on the same probability space. Then for every positive integer m*,

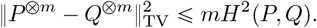

A proof is provided in Appendix C.

In the present setting, we will regard *P* and *Q* two possible gene tree distributions under two distinct species tree parameters, *S*_*P*_ and *S*_*Q*_. Lemma 2 allows us to reduce the problem of determining a lower bound on *m* required to distinguish the two tree parameters into a problem of estimating the Hellinger distance between *P* and *Q*. The general argument is as follows.

Let *f*_max_, *d*_min_ > 0, and assume there exists a continuous function Ψ : (0, *f*_max_] × [*d*_min_, 8) → ℝ_>0_ such that

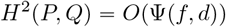

as (*f, d*) → (0, 8). Lemma 2 implies that if

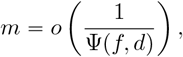

Then

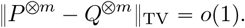

This implies that for any δ > 0 there exists a constant *c*_δ_ > 0 sufficiently small such that

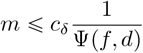

Implies

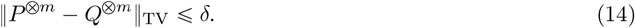

In particular, Eq. (14) precludes accurate species tree estimation when δ is small because it implies that there is no statistical test utilizing only at most 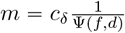 samples whose type-I and type-II error probabilities are both less than 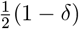. The conclusion that follows is that if

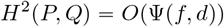

as (*f, d*) → (0, 8), then any hypothesis test to distinguish *S*_*P*_ from *S*_*Q*_ using *m* i.i.d. samples requires at least

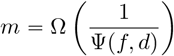

samples to succeed with high probability.

## 3 Distinguishing 2-leaf species trees

### 3.1 Problem statement and key definitions

We first consider the simplest case, that of a species tree with only two leaves. Fix *d*_min_ > 0. Let *d* ⩾ *d*_min_ and let 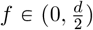.Let *𝒮* be an ultrametric 2-leaf species tree of height *d* with leaves labeled 1 and 2, and define 𝒮 ^′^ similarly, but of height *d* − *f*. The trees *𝒮* and 𝒮 ^′^ are shown in Fig. 2.

**Figure 2:**
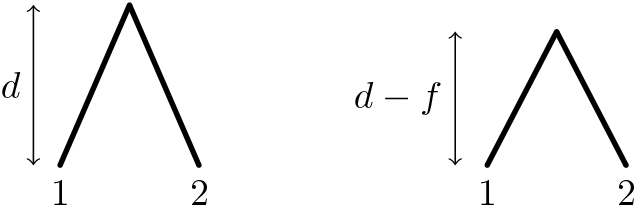
The two tree parameters *𝒮* (left) and *𝒮* ^′^ (right).

Let *θ* be a real-valued random variable with distribution *µ*_*θ*_ such that *µ*_*θ*_ ((*a ∞*))=1, 1 for some *a* 0. For simplicity, write 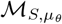 and 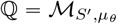.Given data taking the form of *m* i.i.d. gene trees 𝒯_1_, …, 𝒯_*m*_, we wish to distinguish the following two hypotheses:

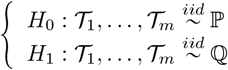

As *f* → 0, these two hypotheses become harder to distinguish.

The distribution of the 2-leaf gene tree 𝒯_1_ is particularly simple, as there is only one coalescence event and it must occur in the root population. Thus, the gene tree can be parameterized by a single variable, namely, the age of the coalescent event *T*. Conditional on *θ, T* has a shifted exponential distribution, with a shift of either *θd* or *θ*(*d −f*) depending on whether the tree parameter is 𝒮 or 𝒮 ^′^; this be verified using Eq. (8).

Let *q*(*t*) and *p*(*t*) denote the density of the gene tree 𝒯_1_ parameterized by the value *T*, under ℚ and ℙ respectively. Using Eqs. (8) and (10),

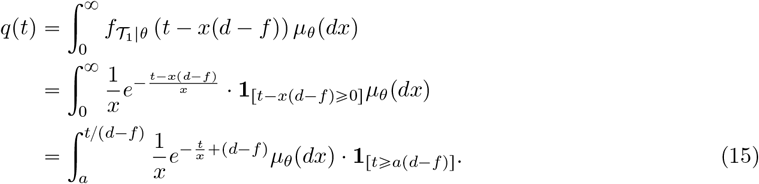

Taking *f* = 0 in Eq. (15) implies

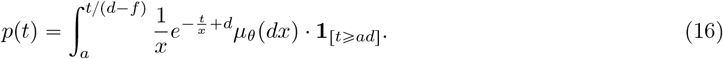

### 3.2 Nonrandom mutation rates

To provide a baseline for comparison with the main result of this section, it is helpful to first consider the case in which genes do *not* exhibit random mutation rates, i.e., the case in which the mutation rate is constant and shared by all genes. In that case, we will show that the number of samples required to distinguish *S*_*d*_ and *S*_*d* − *f*_ with high probability is on the order of *f* ^− 1^.

#### Proposition 1

(Nonrandom mutation rates). *If θ*=*θ*_0_ >0 *is a fixed constant not varying between genes, then*

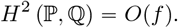

*as* (*f, d*) → (0, ∞), *where the implicit constant in the O*(*f*) *term does not depend on d*.

*Proof of Proposition 1*. Taking *µ*_*θ*_ to be the Dirac point mass at *θ*_0_ and *a* = *θ*_0_ in Eqs. (15) and (16) implies

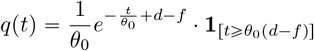

and

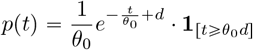

Noting that the intersection of the supports of these two densities is the interval [*θ*_0_*d*, ∞), we have

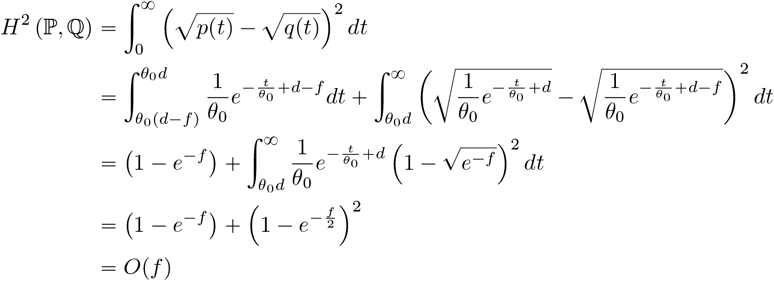

as (*f,d*) → (0, ∞).

By the discussion in Section 2.4.2, we can conclude from Proposition 1 that when mutation rates are constant, distinguishing *S* and *S* ^′^ with high probability as (*f, d*) → (0, ∞), requires that *m* grow at least as fast as *f* ^−1^.

In the proof of Proposition 1, the *O*(*f*) term arises in the region (*θ*_0_(*d* − *f*), *θ*_0_*d*), which is in the support of *q* but not *p*. This suggests that in order to distinguish 𝒮 and 𝒮^′^, one possible strategy is to look for gene trees exhibiting a coalescence age in the interval (*θ*_0_(*d* − *f*), *θ*_0_*d*).Since this cannot occur under *P*, observing such a gene tree would immediately imply that the species tree parameter must be 𝒮^′^, rather than 𝒮. Since the probability of observing such a tree is *O*(*f*) under ℚ, it follows that after *O*(*f* ^−1^) samples, one will have either observed such a tree with high probability (under ℚ), or will be highly confident that such a tree does not exist (under ℙ). Indeed, this approach gives an upper bound of *O*(*f* ^−1^), indicating that the *O*(*f* ^−1^) lower bound from Proposition 1 is tight.

Next, we show that something similar cannot be done when mutation rates are random.

### 3.3 Random mutation rates

In this section, we will establish an information-theoretic lower bound for distinguishing two-leaf species trees when *θ* has a continuous Gamma distribution (*f, d*) → (0, ∞). By **??**, it will be enough to show that

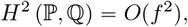

as this will imply that the number of samples required to distinguish 𝒮 and 𝒮^′^, with high probability is of order at least *O*(*f* ^−2^), when *f* is small. To be precise, our first main result is the following, somewhat more general, theorem.

#### Theorem 1

(Lower bound for distinguishing 2-leaf trees). *Let d*_min_ > 0 *be arbitrary. Assume θ has probability density function g with g*(*x*) > 0 *for all x* > 0. *Further assume that there exists a constant C*_gc_ > 0 *such that for all s* > 0, *the following growth condition is satisfied:*

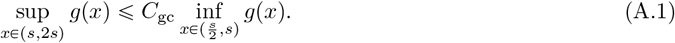

*Then there exists a constant C* > 0 *not depending on f or d such that*

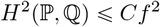

*for all d* > *d*_min_ *and all* 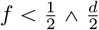.

As a corollary, if *θ* has a Gamma distribution, then it satisfies the assumptions of Theorem 1, and hence the desired conclusion holds.

#### Corollary 1.

*If θ is Gamma distributed then H*^2^(ℙ, ℚ) = *O*(*f* ^2^) *as* (*f, d*) → (0, ∞).

*Proof of Corollary 1*. If *θ* has density 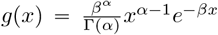 with *α, β* > 0, then clearly *g*(*x*) > 0 for all *x* > 0. It remains only to show that *g* satisfies (A.1), in which case the assumptions of Theorem 1 will be satisfied. To see this, observe that

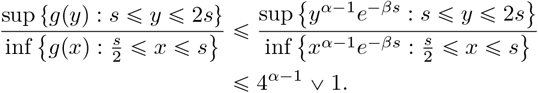

This implies that (A.1) is satisfied with *C*_gc_ = 4^*α*−1^ ⋁ 1.

The proof of Theorem 1 consists of five lemmas, the first of which is nothing but a restatement of the formulas for *p* and *q* established earlier.

#### Lemma 3

(Density formulas for *p* and *q*). *If θ is a nonnegative continuous random variable with density g such that a ≔*inf supp(*g*), *then for all f ∈* [0, *d*),

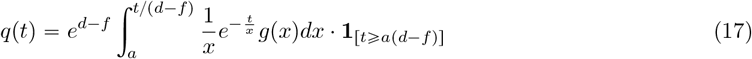

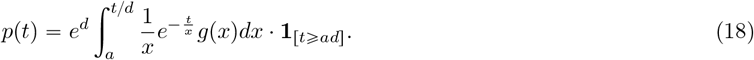

*In particular*, inf supp(*q*) = *a*(*d* − *f*) *and* inf supp(*p*) = *ad*.

*Proof*. Eqs. (17) and (18) follow immediately by Eqs. (15) and (16) with *a* = inf supp (*g*). The claim about the supports of *p* and *q* follows by inspection of these formulas.

The second lemma captures an intermediate estimate which will be used in a later section as well.

#### Lemma 4

(Error term lemma). *Let C*_*E*_ > 0 *and assume that* 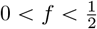.*if*

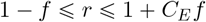

*Then*

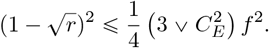

The proof of Lemma 4 is a straightforward application of Taylor’s theorem. The details are given in Appendix D.

The next lemma utilizes Lemma 4 to prove *H*^2^ (ℙ, ℚ) =*O*(*f* ^2^) in the case where *θ* is positive and has a continuous distribution satisfying (A.1).

#### Lemma 5.

*Let θ be a nonnegative random variable with probability density function g such that g*(*x*) > 0 *for all x*> 0. *Further assume that g satisfies* (A.1). *Then there exists a constant C* > 0 *not depending on f or d such that*

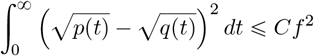

*for all d* ⩾ 1 *and all* 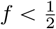.

*Proof of Lemma 5*. Using the assumption that *g*(*t*) > 0 for all *t* > 0, together with the formula for *p*(*t*) in Lemma 3 (with *a* = 0), it follows that *p*(*t*) > 0 for all *t* > 0. Hence

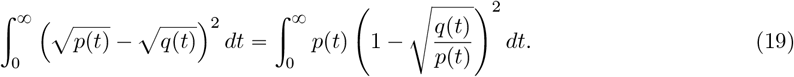

Using the formulas from Lemma 3,

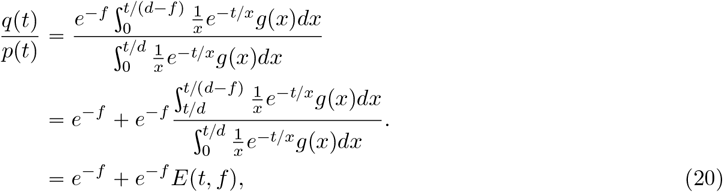

where

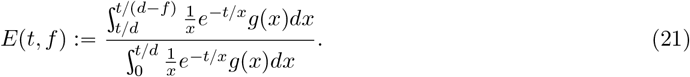

Since *E*(*t, f*) ⩾ 0, it is not difficult to deduce from Eq. (20) that

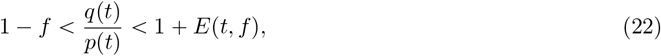

where the first inequality holds because 1 − *f < e*^−*f*^, and the second inequality holds because *e*^−*f*^ < 1. To prove the lemma, it will suffice to show that there exists a constant *C*_*E*_ > 0 such that

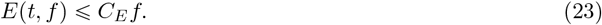

for all *t* > 0, *d* ⩾ 1, and 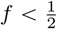, as it will then follow from Eqs. (19) and (22) and Lemma 4 (with 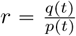) that

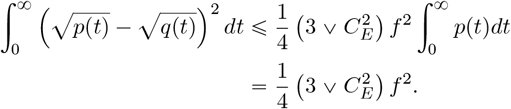

Thus, the rest of the proof will consist of an application of (A.1) to establish the bound in Eq. (23).

By making the domain of integration on the denominator of Eq. (21) smaller,

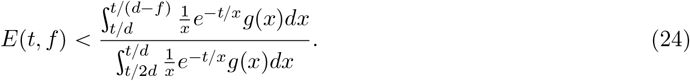

Here we have used the assumption that *g*(*x*) > 0 for all *x* > 0 to ensure that the denominator on the right-hand side of Eq. (24) is nonzero. Because *f* <*d*/2, it follows that

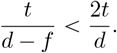

Using this inequality along with the inequality (A.1) (with *s* = *t*/*d*) it follows that

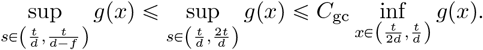

Therefore by Eq. (24)

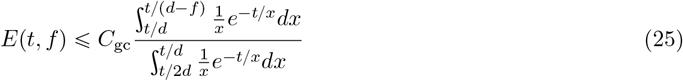

By the substitution 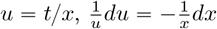,

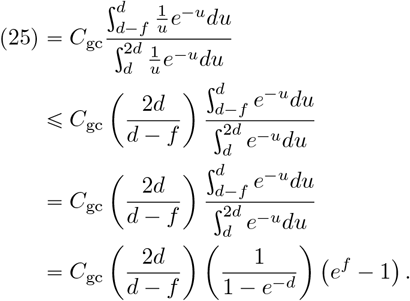

By the assumptions that *d* ⩾ 1 and 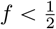, it holds that 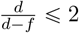 and 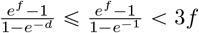 and hence

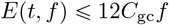

This establishes Eq. (23), completing the proof.

### 3.4 A note on more general mutation distributions

An interesting wrinkle appears when one considers the question of whether Theorem 1 holds under general assumptions about the distribution of *θ*. In Theorem 1 we assumed that *a* ≔ inf supp (*g*) = 0. The alternative, *a* > 0, might arise if *θ* were assumed to be bounded below by a positive constant with probability one. It is plausible, though far from obvious, that knowing this extra information about the mutation rate might allow one to differentiate between ℙ and ℚ with fewer samples than would be required if *a* = 0.

A reasonable heuristic argument suggests that distinguishing 𝒮 from 𝒮 ^‵^ with high probability should still require that *m* grow like *O*(*f* ^−2^), even when *a* ≠0. Suppose that there exists some constant *a* > 0 such that *θ* ⩾ *a* almost surely, and that *θ* has probability density function *g* which is continuous at *a*. Let *S* ∈ {𝒮, 𝒮 ^‵^}. Let 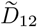 be the coalescent time of 1 and 2 under a gene tree under 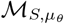. By Lemma 1,

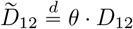

where *D*_12_ is the coalescent time under *ℳ*_*S*,1_.

The essential difference between ℙ and ℚ is that 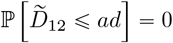, however 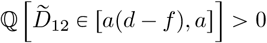. Based on this difference, the obvious strategy to distinguish ℙ and ℚ from *m* independent sampled gene trees is to look for gene trees exhibiting 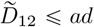.Since this event cannot happen for gene trees sampled from ℙ, we can conclude that if it happens at least once, then the samples must be distributed according to ℚ rather than ℙ. The probability of 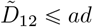 from a single gene tree sampled under ℚ has probability

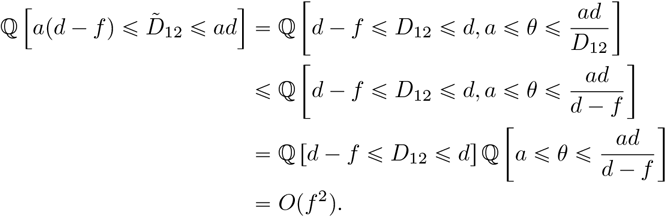

Thus, one requires at least *O*(*f* ^−2^) samples to have a high probability of observing a gene tree with 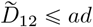 (under ℚ), or to have high certainty that such an event is not possible (under ℙ). This argument suggests that with *O*(*f* ^−2^) samples, one can distinguish the two hypotheses with high probability, and on the basis of this heuristic argument it is reasonable to guess that *requires O*(*f* ^−2^) samples, as we showed in the case where *a*= 0.

Yet surprisingly, the case with *a* >0 does not admit a proof similar to the one used to prove the lower bound for *a* = 0. In particular, we will provide an example in which *a* > 0 and *H*^2^ (ℙ, ℚ) = *ω*(*f*^2^). As we will show, the reason for the difference in qualitative behavior between the *a* = 0 and *a* > 0 cases concerns the magnitude of the integral

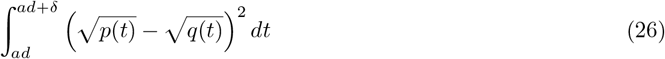

when *δ, d* >0 are fixed and *f* is very small. On one hand, when *a* >0, the formulas from Lemma 3 imply that *p*(*ad*) = 0 and *p*(*ad*) >0 (see, e.g., Fig. 3), and this can cause the size of the integrand to spike when *t* is just slightly larger than *ad* (see Fig. 4). Moreover, this spike may not decrease quickly enough for the above integral to be *O*(*f* ^2^). Since Eq. (26) is a lower bound for *H*^2^(ℙ, ℚ), it follows that *H*^2^(ℙ, ℚ) ≠ *O*(*f* ^2^). On the other hand, this situation cannot arise when *a* = 0, since in that case Lemma 3 implies that *p*(*ad*) = *q*(*ad*) = 0.

**Figure 3:**
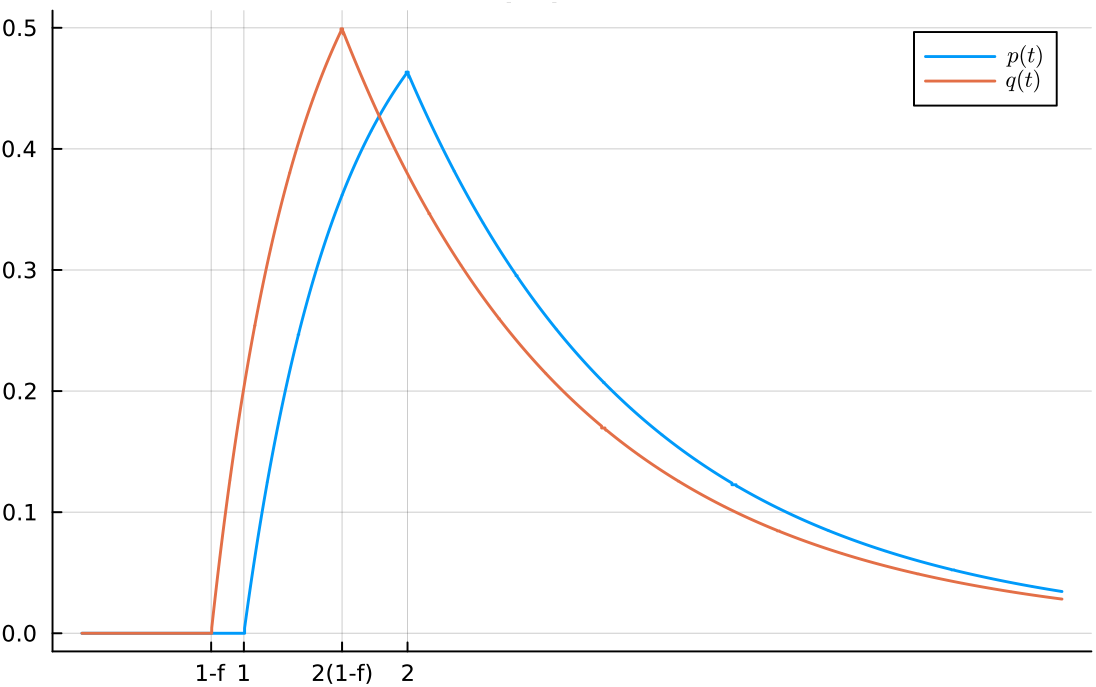
A plot of *p* and *q* when *g*(*t*) = **1**_[1,2]_(*t*) and *f* = 0.2.

**Figure 4:**
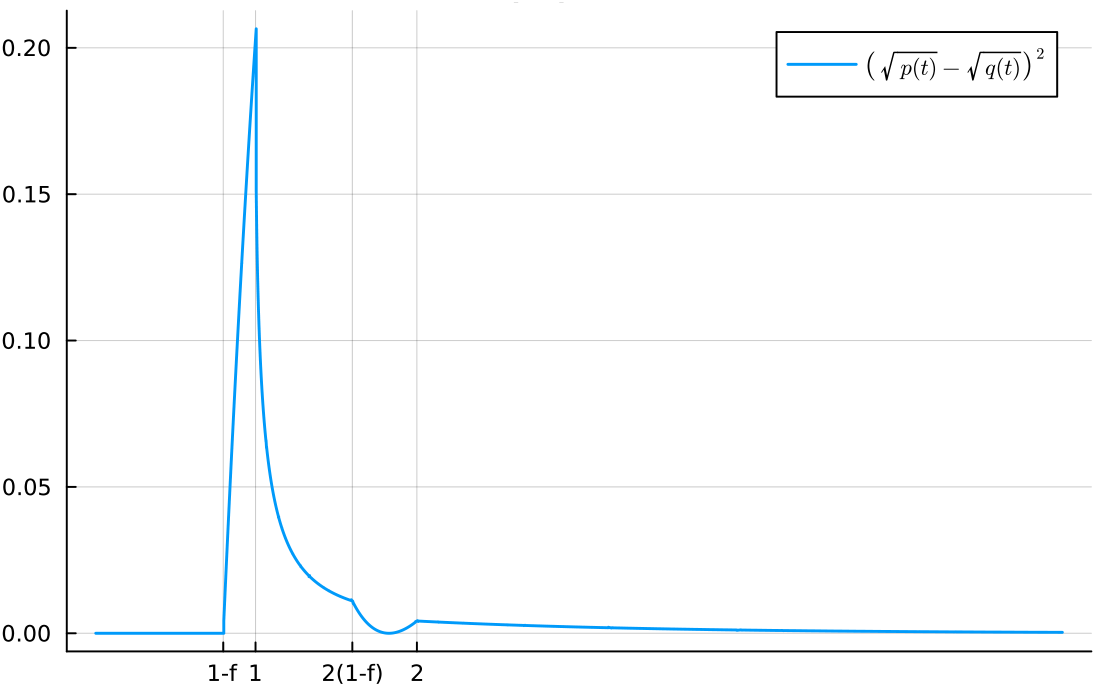
A plot of the function 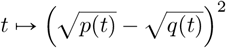 when *g*(*t*) =**1**_[1,2]_ (*t*) and *f* = 0.2.

**Figure 5:**
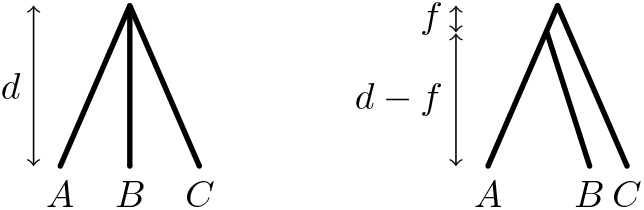
The two tree parameters 𝒮_0_ (left) and 𝒮_*f*_ (right).

The specific example we will consider is the case where *θ* is uniformly distributed on the interval r1, 2s, so that *g*(*t*) = **1**_[1,2]_(*t*) and *a* = 1. For simplicity, in the remainder of this section we will assume that *d* = 1. We will show that for *g*(*t*) = **1**_[1,2]_, the the conclusion of Theorem 1 does not hold. This is surprising given that *g* is a nothing more than a translation of **1**_[0,1]_, a function which satisfies (A.1). The graphs of *p* and *q* for this choice of *g* are shown in Fig. 3; we note that *p* (1) 0 but *q* (1) > 0, and the resulting spike can be seen in Fig. 4.

We begin by taking the first-order Taylor expansion of *p* and *q* at the point (*t, f*) = (1, 0).

#### Lemma 6.

*Regarding q and p as functions of both t and f, it holds that*

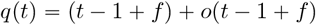

*and*

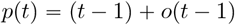

*as* (*t, f*) → (1, 0) *with t* ⩾ 1 *and f* ∈ [0, 1).

*Proof of Lemma 6*. We show only the result for *q*(*t*), since the result for *p*(*t*) follows from the same calculuation but with *f* = 0.

Let 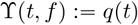, where we regard *q* as a function of both *t* and *f*, and which is defined on for all *t* ⩾ 0 and all *f* ∈ [0, 1). By Lemma 3,

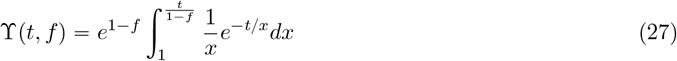

for all *t* ⩾ 1. Making the substitution 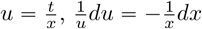 gives

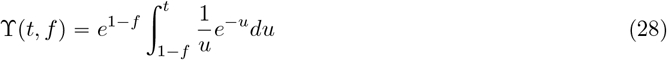

and hence

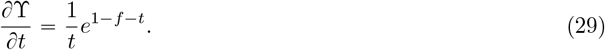

Next we compute 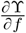.Using the product rule and the Leibniz integral rule to differentiate both sides of Eq. (27) with respect to *f* :

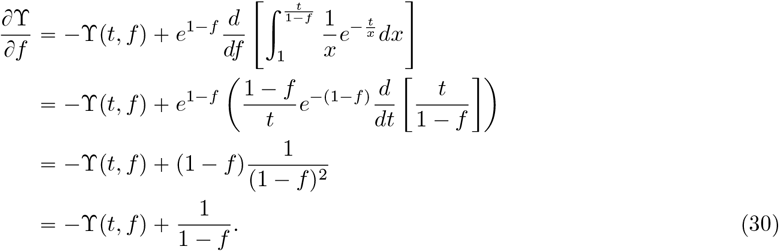

Using Eqs. (28) to (30), we deduce that 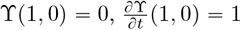, and 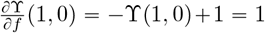. Therefore first-order Taylor expansion of *q* about the point (*t, f*) = (1, 0) is

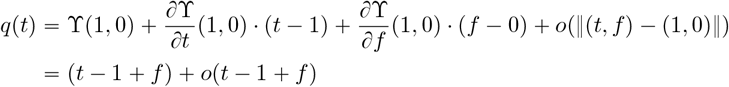

as (*t, f*) → (1, 0) with *t* ⩾ 1 and *f* ⩾ 0.

By Lemma 6, *p* and *q* satisfy the hypotheses of the following lemma, which says that the integrand 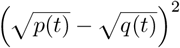 has too much mass near *a* for its integral to be *O*(*f* ^2^). A picture of this is shown for our specific example in Fig. 4.

#### Lemma 7.

*Assume that*

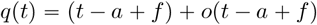

*and*

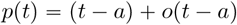

*as* (*t, f*) → (*a*, 0) *with t* ⩾ *a and f* ∈ [0, *d*). *Then*

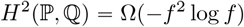

*Proof of Lemma 7*. By assumption,*q* = (*t* − *a* + *f*) + *ϵ*_*q*_(*t, f*) for some function *ϵ*_*q*_ = *o*(*t* − *a* + *f*). Then

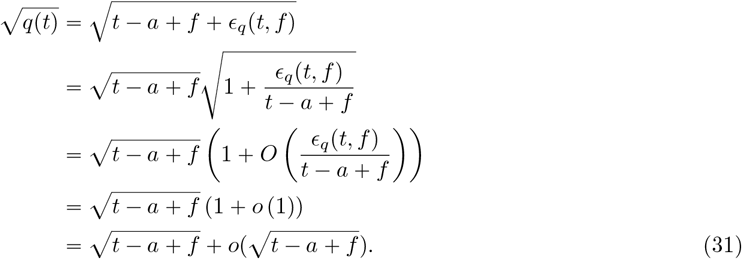

Taking *f* = 0 in the above calculation gives

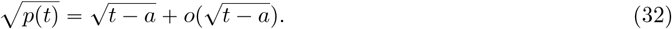

By Eqs. (31) and (32)

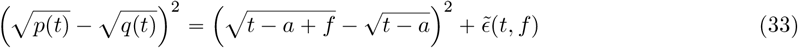

where 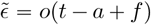. Therefore there exists δ*>* 0 such that 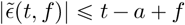 whenever |*t− a* |*+ f*|< *δ*. This, in turn, implies that

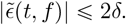

Combining this with Eq. (33) implies that

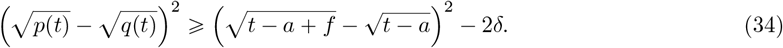

for all *t* (∈*a, a* +*δ*) and all 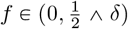. Therefore

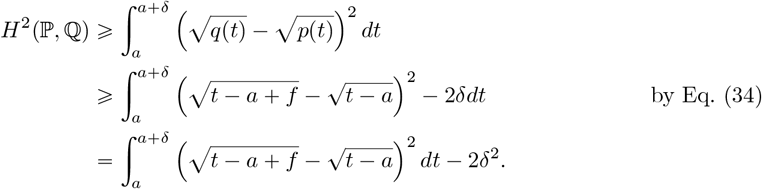

Making the substitution *t* ↦ *t* +*a*,

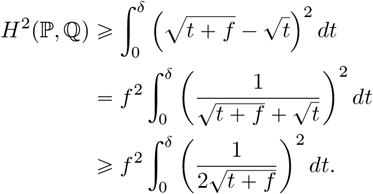

Evaluating the integral on the right hand side gives

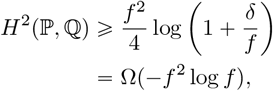

which completes the proof.

## 4 Distinguishing a binary 3-leaf tree from a star tree

In this section we consider the problem of distinguishing 3-leaf species trees using gene trees drawn from the multispecies coalescent, assuming mutation rates are gamma-distributed with an atom at zero.

### 4.1 Statement of main result

For each *l* 0, let 𝒮_*l*_ be a binary tree with three leaves *A, B* and *C*, rooted binary tree topology *AB* |*C* such that the internal branch length, in coalescent units, is *l* > 0 and the age of the root is *d*> *l*. Let *AB* denote the parent population to *A* and *B*, and *ABC* the root population.

**Problem:** Given *m* i.d.d. gene trees 𝒯_1_, …, *𝒯*_*m*_ drawn from the MSC with random mutation rates, the hypothesis testing problem is to distinguish between the following two hypotheses

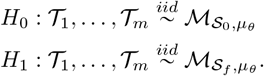

Our main result is the following theorem, which shows that in order to succeed with high probability as (*f, d*) → (0, ∞), it is necessary that *m* grow like *f* ^−2^.

#### Theorem 2

(Main result). *Suppose θ has a gamma distribution µ*_*θ*_, *possibly with an atom at zero. Then for every ϵ* > 0 *there is a constant c such that*

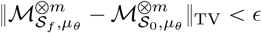

*whenever*

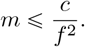

*Moreover, c does not depend on d, m, or f, but may depend on the shape parameter of the Gamma distribution*.

Before proving Theorem 2, we first consider the special case where *µ*_*θ*_ is a gamma distribution with no atom at zero. In particular, the majority of the work will be in proving the following proposition, which will then be used to prove the more general result of Theorem 2:

#### Proposition 2

(Gamma-distributed mutation rates). *Fix* 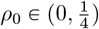.*Assume that θ has Gamma-distribution µ*_*θ*_ *with parameters α, β. Then there exists a constant C* > 0, *depending only on α, such that*

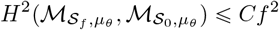

*whenever* 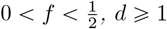,*and* 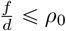.

### 4.2 The density functions 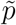 and 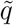

With probability one, the topology of a 3-leaf gene tree 𝒯 under the MSC exhibits one of three possible topologies: *AB*| *C, AC* |*B*, or *BC* |*A*. Let *K∈* {0, 1, 2, 3} denote the topology of 𝒯, with *K* 0 corresponding to the star topology, *K* = 1 corresponding to *AB C*, and *K* = 2 and *K* = 3 corresponding to the other two binary tree topologies. Thus, we can represent a gene tree drawn according to the MSC by a triple *𝒯* = (*K, T, S*), where

- *K* ∈ (0, 1, 2, 3) indicates the rooted tree topology of *𝒯*,
- *T* is the time until first coalescence, and
- *S* is the waiting time between the first and second coalescence events.

Parametrized in this way as a triplet 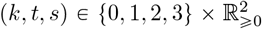, we can identify the space of ultrametric 3-leaf gene trees with the measurable space

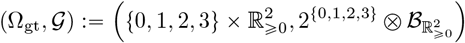

where 2^{0,1,2,3}^ is the power set of {0, 1, 2, 3} and 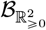 is the standard Borel *σ*-algebra on 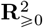.

Define a measure *µ*_dom_ on Ω_gt_ by *µ*_dom_ ≔ (*δ*_1_ + *δ*_2_ + *δ*_3_) ⊗ λ_L_ ⊗ λ_L_ where *δ*_*i*_ denotes the Dirac measure on *i* ∈ {1, 2, 3}, and λ_L_ denotes the Lebesgue measure on ℝ _⩾0_. Then for any measurable *A* ∈ 𝒢,

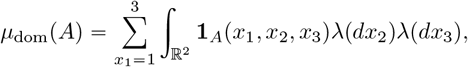

#### Remark 1

(Key notation). *Let* 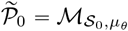 *and* 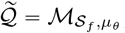, *where µ*_*θ*_ *is gamma-distributed with density function g. In words*, 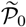 *is the gene tree distribution under the MSC on species tree 𝒮*_0_ *with random mutation rate θ* ∼ *µ*_*θ*_ *and* 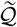 *is similar but with tree 𝒮*_*f*_. *Further, let* 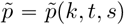 *and* 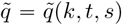 *denote the probability density functions of a gene tree 𝒯* = (*K, T, S*) *under* 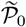 *and* 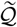 *respectively, with respect to the dominating measure µ*_dom_.

The next proposition gives explicit formulas for 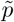 and 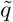.

#### Lemma 8

(Scaled density formulas). *Let f* ∈ [0, 1) *and d* > *f, and assume that θ is a positive random variable with probability density function g. Then*

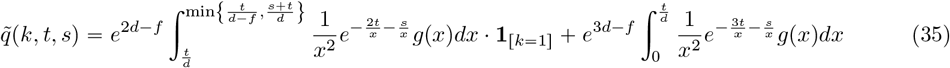

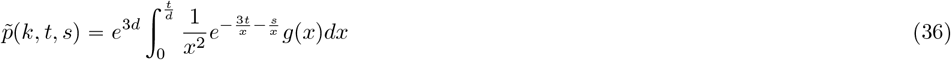

*for all s, t* > 0 *and all k* ∈ {1, 2, 3}. *For k* = 0, *we have* 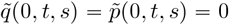 *for all t, s* ⩾ 0.

*Proof of Lemma 8*. It will be enough to compute 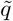, as the computation for 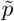 follows by the same argument with *f* = 0. Let *x* > 0 be a real number. We begin by computing the conditional distribution of a gene tree given *θ* = *x*; that is, suppose 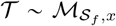.Since *x >* 0, it follows that 𝒯 has a binary tree topology with probability one, so 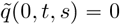 for all *t, s* ⩾ 0 and without loss of generality we may assume that *k* ∈ {1, 2, 3} in the remainder of this proof.

Let 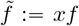 and 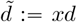. Since > is obtained by running the multispecies coalescent on 𝒮_*f*_, there are two disjoint cases, illustrated in Fig. 6. Either:

**Figure 6:**
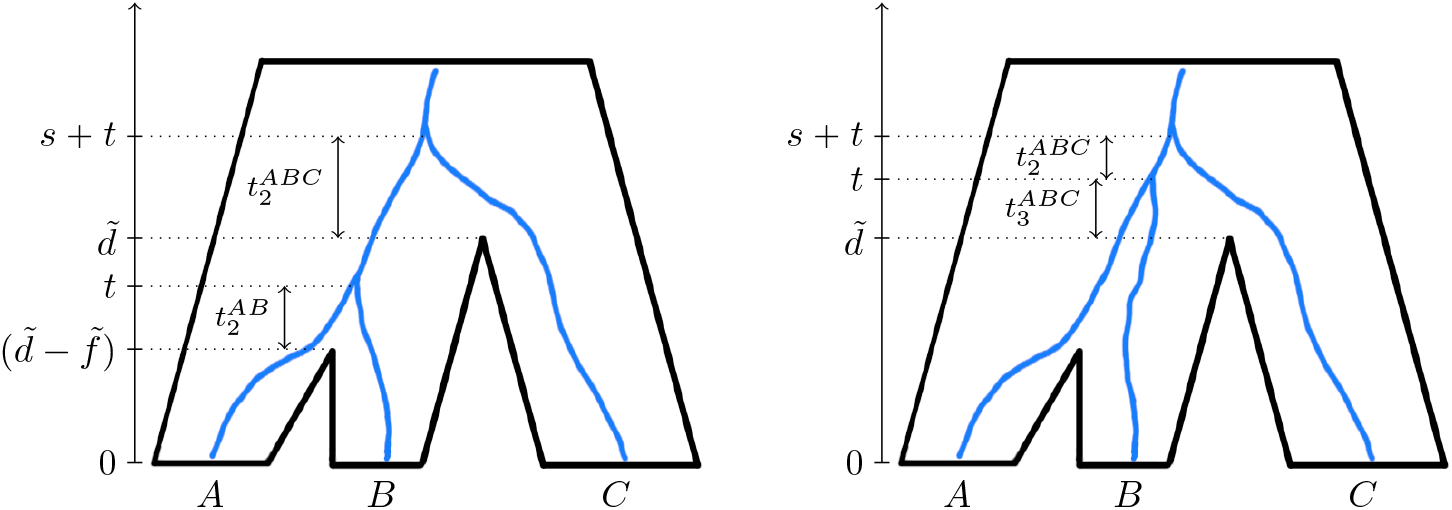
A depiction of the two disjoint cases for a 3-leaf gene tree. *Left:* a gene tree whose first coalescence event occurs below the root. Here, 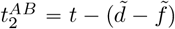 and 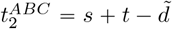.*Right:* a gene tree whose first coalescence occuring above the root, so that 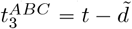 and 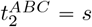.

i. the first coalescence occurs *below* the root, in which case *k* = 1 with probability one, or
ii. the first coalescence occurs *above* the root, in which case the three topologies *AB* |*C, AC* |*B*, and *BC* |*A* are equally likely to be exhibited by the gene tree.

Using the above observations along with Eq. (8), the mixed density of *𝒯* is given by

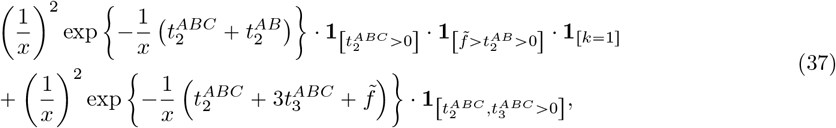

where the first term corresponds to case (i.) and the second term corresponds to (ii.). In particular, 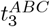 is undefined in case (i.), so by convention we take this to mean that 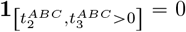, and hence the second term vanishes in case (i.). On the other hand, *t*^*AB*^ is undefined in case (ii.), so 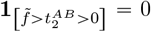, and hence the first term vanishes in case (ii.)

Let *t* be the time of the first coalescence event and let *s* be the waiting time between the first and second coalescent events, both measured in expected number of mutations per site. It is easy to see from Fig. 6 that under case (i.) we have

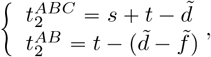

whereas under case (ii.),

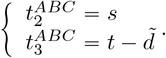

Therefore we can write Eq. (37) as a function of *t, s* and *k*. After simplification, the density function of 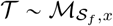 is

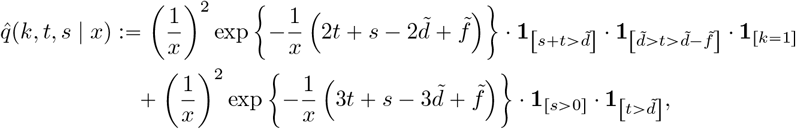

and since 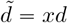 and 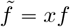,

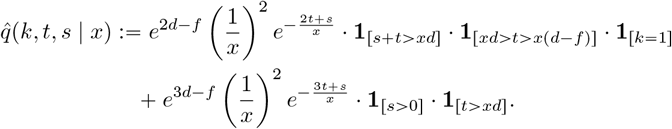

Now let 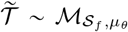, where *θ* is a positive random variable with probability density function *g*. By Eq. (10), the mixed density of 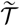 is

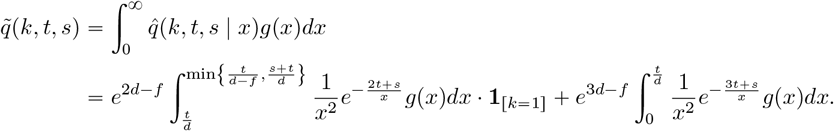

The analogous formula for 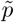 is obtained by taking *f* = 0 in the above formula.

### 4.3 Proof of Proposition 2

Fix *f*_0_ ∈ (0, 1/2) and assume 0 < *f* < *f*_0_. Since 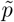 and 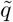 are both absolutely continuous with respect to *μ*_dom_, the Hellinger distance can be expressed as

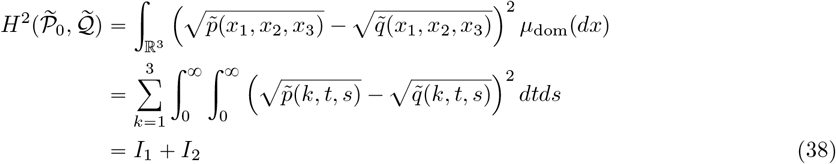

where

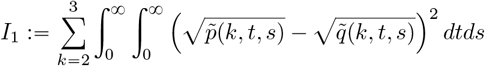

and

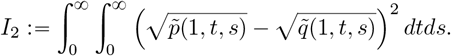

We will show that both *I*_1_ and *I*_2_ are both *O*(*f* ^2^).

When *k =* 2 or *k =* 3, Lemma 8 implies that 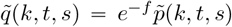. This observation allows *I*_1_ to computed exactly, so an estimate is readily obtained, as shown in the next lemma.

#### Lemma 9

(Estimate for *I*_1_). *For all f* > 0 *and all d* ≥1, *it holds that*

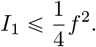

*Proof of Lemma 9*. By Eqs. (35) and (36), we have

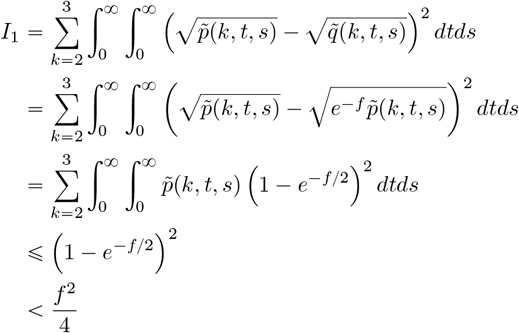

where we used the numerical inequality 1 − *e*^−*x*^⩽ *x* with 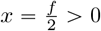.

It remains to show that *I*_2_ = *O*(*f* ^2^). The estimation of *I*_2_ is more involved, and we split our analysis into two cases: *s* ⩽ 10*t* and *s* > 10*t*. That is, let 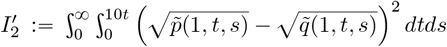 and let 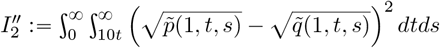 so that

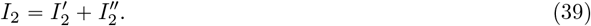

We will bound 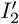 and 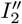 indidivually.

#### 4.3.1 Estimate of 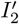 (i.e., when *s* ⩽10*t*)

The next two lemmas estimate the portion of the integral *I*_2_ with 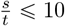. For this case, the estimate can be established in a manner similar to the proof of Lemma 5, i.e., by Taylor expanding 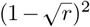 as in Lemma 4, and showing that

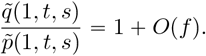

We begin with the following technical lemma.

##### Lemma 10.

*Assume f* ⩽ 1/2 *and d* ⩽ 1. *Let* 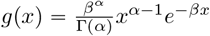 *where α, β* > 0, *and let*

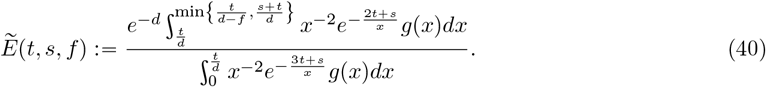

*Then for all s, t* > 0 *such that* 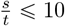, *the following estimate holds:*

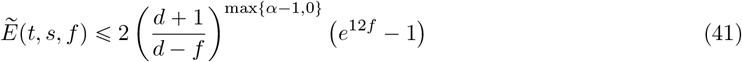

*In particular, this implies that for all f* ⩽ 1/2 *and all d* ⩾ 1,

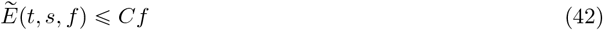

*where C* =1610 · 4^max{*α*−1,0}^

*Proof*. Making the domain of integration on the numerator of Eq. (40) possibly larger, and using 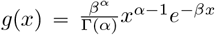, we have

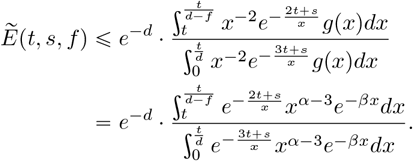

Since *e*^−*βx*^ <1, replacing *e*^−*βx*^ by 1 in both the numerator and denominator makes the right-hand side larger, so

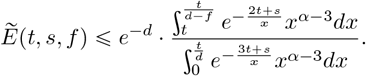

Making the substitution 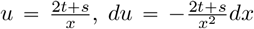 in the numerator and 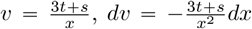 in the denominator yields

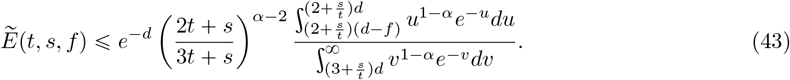

There are two cases, depending on whether *α* is greater or less than 1:

- If 0 < *α* ⩽ 1, then

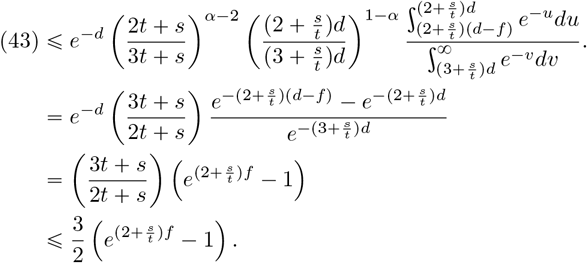 Therefore by the assumption 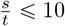,

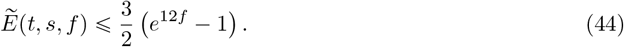
- If *α* > 1, then making the domain of integration on the denominator smaller in Eq. (43) implies

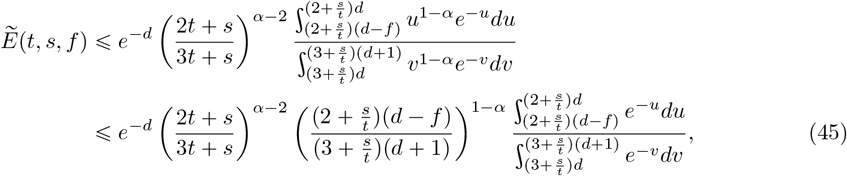

where, in going from the first to the second line, we have utilized the assumption that *α* > 1 to bound the term *u*^1−*α*^ above by replacing *u* with its lower integration bound, and bound the term *υ*^1−*α*^ below by replacing *υ* by its upper integration bound. Next, we simplify the constants and evaluate the integrals:

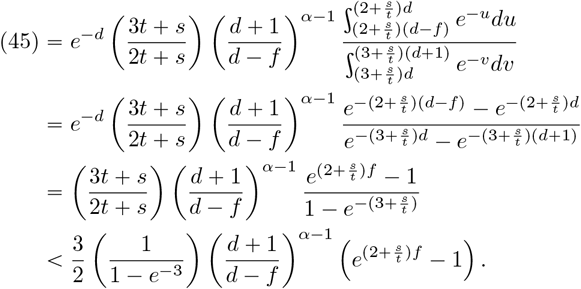

Using the assumption that 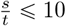 and the observation that 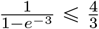, it follows that

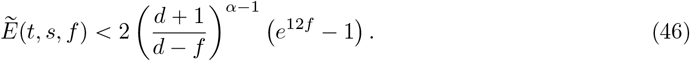

Taken together, Eqs. (44) and (46) imply Eq. (41) in the statement of the lemma. Finally, Eq. (42) can be deduced from Eq. (41) by considering the worst case (*f, d*) = (1/2,1).

The next lemma applies the estimate from Lemma 10 to establish a bound on 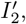 i.e., is the case where *s* ⩽10*t*.

##### Lemma 11

(Estimate for *s* ⩽10*t*). *There is a constant C* > 0 *not depending on f or d such that*

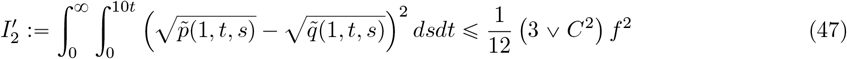

*whenever* 0 <*f* ⩽ 1{2 *and d* ⩾1.

*Proof of Lemma 11*. The proof of this lemma is similar to that of Lemma 5. By the formula for 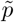 from Lemma 8, it it easy to see that 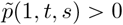 for all *t, s* > 0, and hence

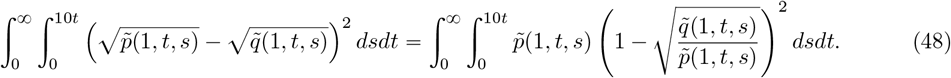

Using the formulas from Eqs. (35) and (36), we can write

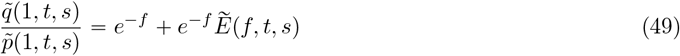

where 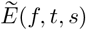 is defined in Eq. (40). Since 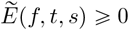 and 1 − *f* < *e* ^−*f*^ < 1, it holds that

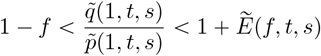

By the bound established in Lemma 10, there exists a constant *C* >0 not depending on *d* ⩾ 1 or *f* ⩽ 1/2 such that

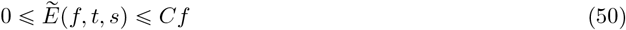

for all *s, t* > 0, such that 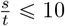. Therefore it follows from Eq. (48) and Lemma 4 that

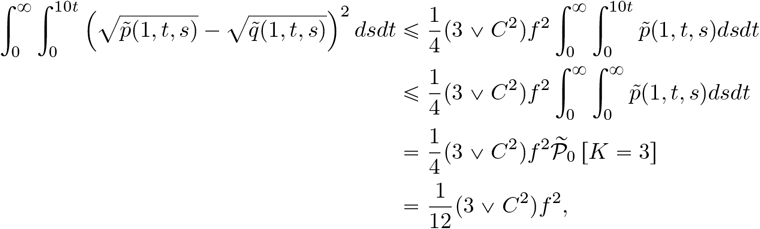

where the last equality follows immediately by symmetry: since 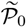 is the distribution of a 3-leaf gene tree under the MSC on parameterized by a star tree, all three rooted binary tree topologies are equally likely.

#### 4.3.2 Estimate of 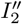 (i.e., when *s* ⩾ 10*t*)

The next lemma handles the case wehere 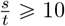.

##### Lemma 12

(Estimate for *s* ⩾ 10*t*). *Let* 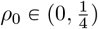. *There exists a constant C* >0 *not depending on d or f such that*

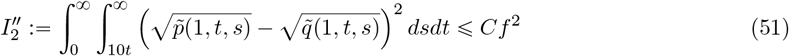

*for all* 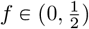 *and d* ⩾ 1 *such that* 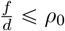.

*Proof of Lemma 12*. For each *u* ⩾ 0 and *s, t* > 0 such that 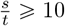, let

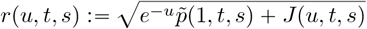

and let

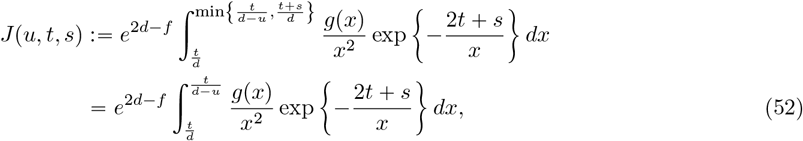

where the second equality holds since *s* > *t* implies that 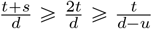 for all *u* ⩽*f* ⩽*d*/2.

Observe that function *u* ⟼ *r*(*u, t, s*) is differentiable on the interval[0, *f*] with continuous partial derivative

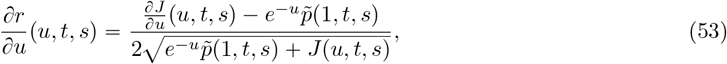

where

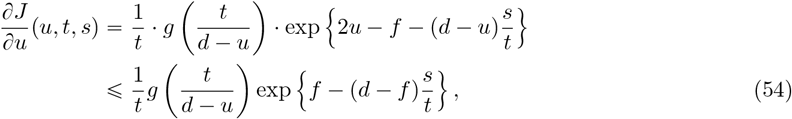

where the equality follows by an application of the Leibniz differentiation rule, and inequality follows trivially by the assumption 0 < *u* ⩽*f*.

In addition, for all *s* >*t* > 0, the function *r* satisfies the boundary conditions

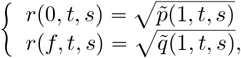

so that by the fundamental theorem of calculus,

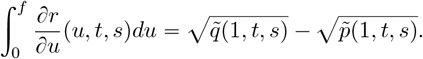

Therefore, using the Minkowski inequality for integrals,

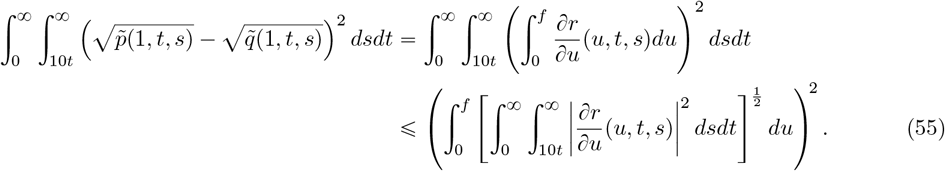

Therefore to prove Eq. (51), it will suffice to show that there exists a constant *C* >0 not depending on *f, d* or *u* such that

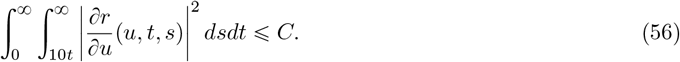

We begin by bounding the integrand of Eq. (56). Let

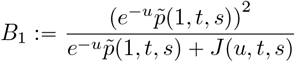

and

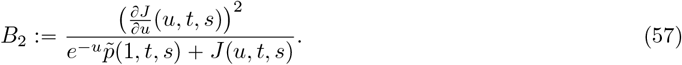

Using Eq. (53) as well as the numerical inequality (*x* ^2^ − *y*^2^) ⩽ *x*^2^ + *y*^2^, (*x, y* ⩾0),

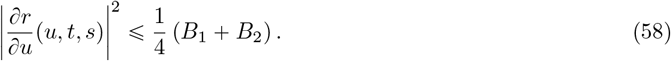

We will bound *B*_1_ and *B*_2_ individually.

Since *u* ⩾ 0 and *J*(*u, t, s*) ⩾0 for all *u* ⩾ 0 and *t, s* > 0, and since 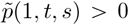 for all *s, t* >0, the quantity *B*_1_ admits the following bound:

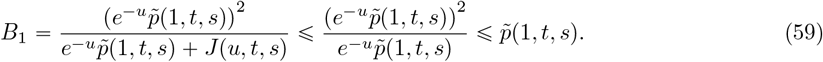

Next we will bound *B*_2_. By the inequalities *J*(*u, t, s*) ⩾ 0, and *u* ⩾*f*, it follows trivially that

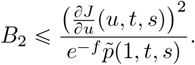

By Eq. (54),

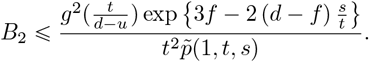

Plugging in the formula for 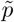 from Lemma 8,

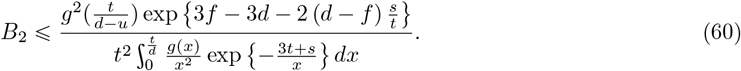

Let

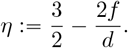

By the assumption that 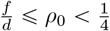,

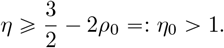

In particular, since *η* > 1, it holds that 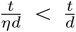. Therefore the right-hand side of the Eq. (60) can be made larger by integrating over the smaller domain 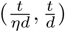. Doing this, and then substituting in the gamma desnity 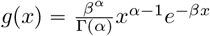, yields

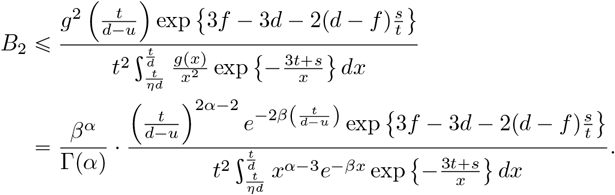

Next, we will bound the numerator above using the inequality 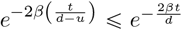, and bound the denominator below using the inequality 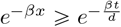, which holds for all 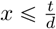. This gives the inequality

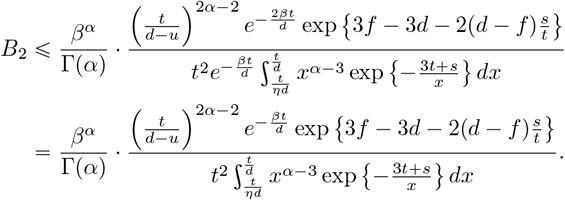

Observe that for all 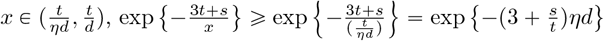. Therefore

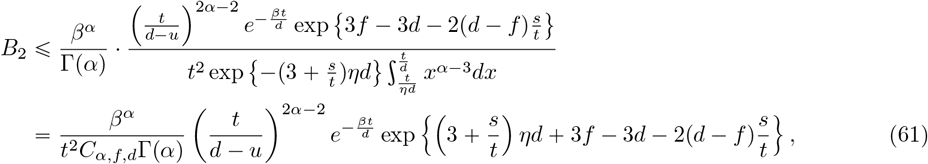

where

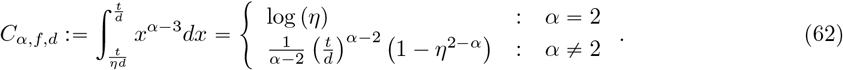

To bound Eq. (61), we shall use the assumption that *s* ⩾ 10*t*. Since 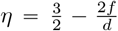, the quantity in curly brackets in Eq. (61) can estimated as follows:

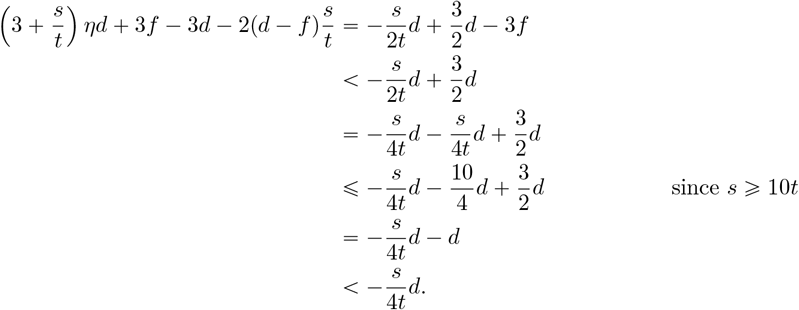

Plugging this inequality into Eq. (61) yields

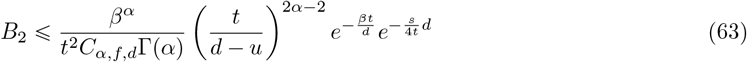

for all *s, t* > 0 with *s* ⩾ 10*t*. Next, we will use Eq. (63) to bound the integral 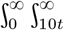 *B*_2_(*u, t, s*)*dsdt* above by a universal constant not depending on *f, d* or *u*; however, the calculations differ somewhat depending on whether *α* = 2 or *α* ≠ 2, so we consider these two cases separately.

##### Case 1.

Suppose *α =* 2. Plugging in the definition of *C*_*α,f,d*_ from Eq. (62) into the right-hand side of Eq. (63) and simplifying gives

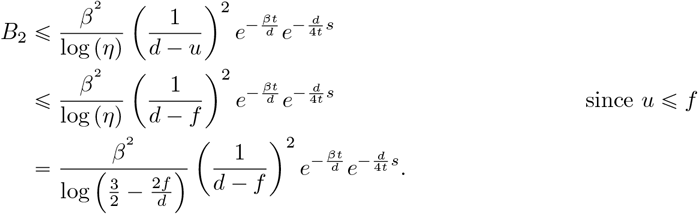

Integrating both sides,

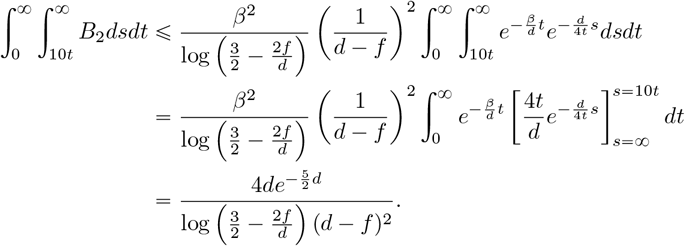

Using the assumptions *d* ⩾ 1 and 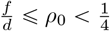 to bound the right-hand side, it follows that that there exists a constant *C*_*α*=2_ > 0 not depending on *f* or *d* such that

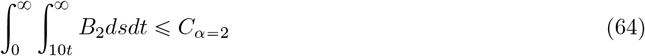

for all 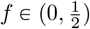 and *d* ⩾ 1 satisfying 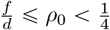.

##### Case 2.

Suppose *α* ≠2. Plugging in the definition of *C*_*α,f,d*_ from Eq. (62) into the right hand side of Eq. (63) and simplifying gives

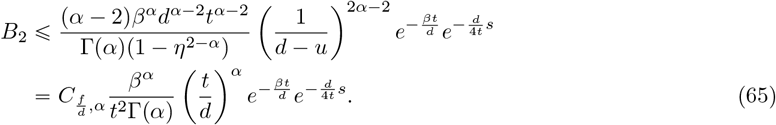

where

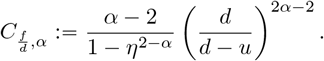

We note that 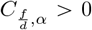 for all *α* ≠ 2. Moreover, using the inequalities 0 ⩽ *u* ⩽ *f* and 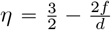, we can deduce that

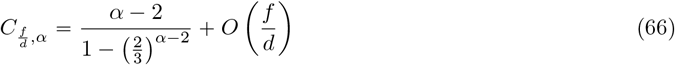

as 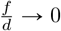.

Integrating both sides of Eq. (65) gives

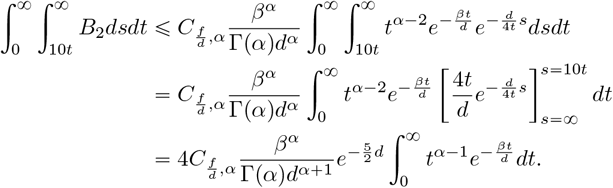

Using properties of the Gamma distribution,

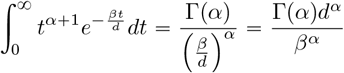

Plugging this into the previous equation gives

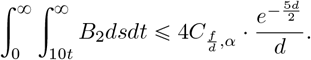

Together with Eq. (66), this implies that if *α* ≠ 2 then exists a constant *C*_*α*≠2_ > 0 not depending on *f* or *d* such that

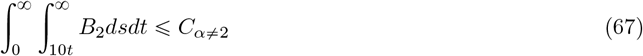

for all 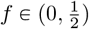 and *d* ⩾ 1 with 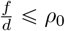. This completes the estimate for Case 2.

Putting the two cases *α* = 2 and *α* ≠ 2 together, we conclude from Eqs. (64) and (67) that there exists a constant 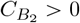 not depending on *f* or *d* such that

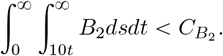

for all 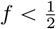 and *d* ⩾ 1 such that 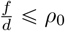. Therefore by Eqs. (58) and (59),

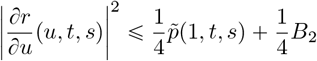

and therefore

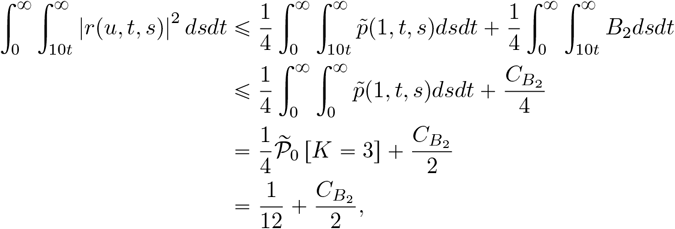

where the last equality follows immediately by symmetry: since 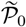 is the distribution of a 3-leaf gene tree under the MSC on parameterized by a star tree, all three rooted binary tree topologies are equally likely. This proves Eq. (56), and hence it follows by Eq. (55) that

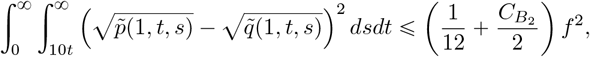

which completes the proof of the lemma.

#### 4.3.3 Final analysis for proof of Proposition 2

Recall that we defined 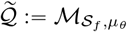 and 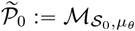. Using Eqs. (38) and (39),

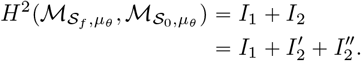

Plugging in the estimates for *I*_1_, 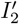 and 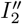 from Lemmas 9, 11 and 12 implies the statement of Proposition 2.

### 4.4 Proof of Theorem 2

We are now ready to prove Theorem 2 and Appendix C. The main idea of the proof is to use the elementary properties of total variation distance discussed in Section 2.4.1 to reduce the problem to the case of gamma-distributed mutation rates with no atom at zero, allowing us to apply the bound from Proposition 2.

*Proof*. Throughout this proof, we will use the notation *μ*_*W*_ to denote the probability measure induced by the random variable *W*. By assumption we have 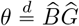 where

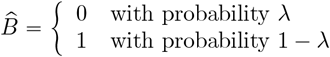

for some λ ∈[0,1), and 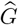 is an independent gamma-distributed random variable with shape and rate parameters *α, β* > 0. Hence

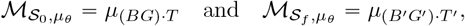

where 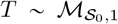 and 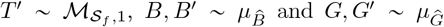, where 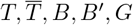, and *G*^1^ are all independent. Hence, we need to bound the total variation distance between 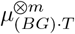 and 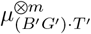.

Let (Ω_gt_, 𝒢) be the space of ultrametric 3-leaf gene trees as defined in Section 4.1, and define 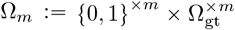 and 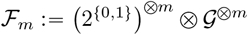.

Let 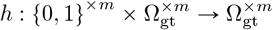 be the map given by

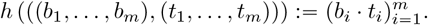

Then *h* is a measurable function (see Lemma 15 in Appendix E for proof).

We’ll need to introduce some variables and notation. 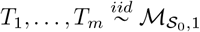 and 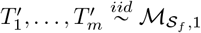, let 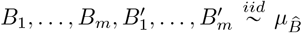 and let 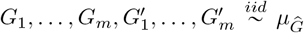. Further, define 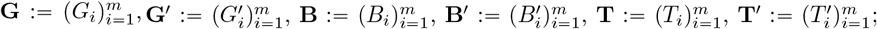 these six random vectors are all assumed to be independent. Finally, define 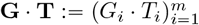 and 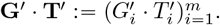.

Observe that *h* (**B, G · T**) and *h* (**B**^1^, **G**^1^ · **T**^1^) have distributions 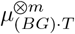 and 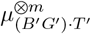, respectively. By Lemma 13, the mapping *h* can not increase the total variation distance, and hence

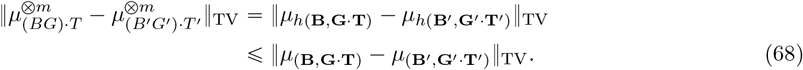

Moreover, by the independence properties **B** ╨ **G** · **T** and **B**^1^ ╨ **G**^1^ · **T**^1^, the meausures *μ*_(**B**,**G**_·_**T**)_ and 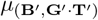 are both product measures on (Ω_*m*_, *F*_*m*_), and hence by Lemma 14,

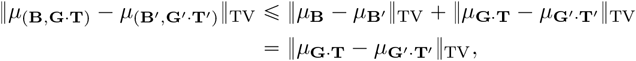

where the equality follows trivially because **B** and **B**^1^ have the same distribution. Combining the above inequality with Eq. (68), we obtain

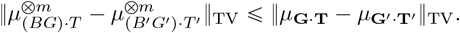

Since 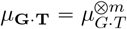 and 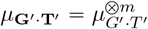, the right-hand side equals 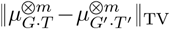, and therefore Lemma 2 implies

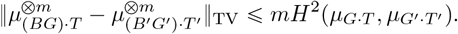

Using Proposition 2 to bound the right-hand side, it follows that for any 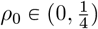, there exists a constant *C* >0 depending only on *α* such that

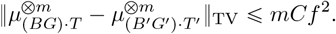

for all 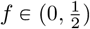, *d* ⩾ 1 with 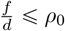. We conclude that if *m =o (f* ^−2^) as (*f, d*) ⟶ (0 ∞), then

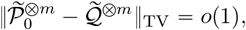

which implies the statement of the theorem.

## 5. Discussion

In Appendix A, we give an example in which a 3 leaf tree *S* with topology 12|3 could be distinguished from a 3-leaf star tree *S*^⋆^. In that case, the assumption of constant mutation rate ensured that presence of ILS on a gene tree could be detected from the branch lengths of the gene trees, and it was this fact that allowed high estimation accuracy to be achieved with as low as *m* = *O*(*f*) samples. In Theorem 2, we showed that such performance cannot be replicated with randomly-varying mutation rates. In particular, Theorem 2 implies that as *f* ⟶0, consistent estimation of the species tree requires that the number of samples satisfy

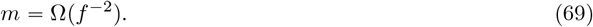

This is a substantially higher data requirement when *f* is small. Moreover, this lower bound is tight, since it is matched by the *m* = *O*(*f*^−2^) upper bounds achieved by topological inference methods—i.e., methods which make no use of branch lengths at all, as shown in Fig. 1. One implication of this is that, for data exhibiting mutation rate variation, inference methods utilizing information about gene tree branch lengths may be unlikely to substantially outperform methods which do not utilize such information.

We conclude with some directions for future research. First, our results hold when mutation rates follow a continuous gamma distribution, possibly with an atom at zero. In the case that mutation rates follow a *discrete* gamma distribution [5], consistent estimation may be possible using fewer than *O*(*f*^−2^)genes; it would be of interest to establish lower or upper bounds for that case as well, as the discrete gamma distribution is also widely used in phylogenetic analyses.

Second, in this study we assumed that data takes the form of *m* error-free gene trees. In the setting where gene trees are estimated via gene sequences, we expect the lower bound to be at least as large as that given in Eq. (69), since in that case one has only imperfect information about the gene trees, a fact which can only make the estimation problem harder. However, in the case of gene sequence data, we also expect the data requirement to depend nontrivially on *d*.

Finally, we caution that it is not at all obvious whether Theorem 2 generalizes to trees with more than three leaves. Indeed, it is conceivable that for large trees (i.e., as *n*⟶ ∞), there may exist some method of estimating the mutation rates corresponding to each gene tree, and then using those estimates to achieve consistent estimation of the species tree topology with *m* = *O*(*f*^−2^)samples. Finding such a method—or showing that no such method exists—would help to further clarify the effect of variable mutation rates on the sample complexity of species tree estimation. See, e.g., [37, 38] for some related results.

## Acknowledgements

This paper is based upon work supported by the NSF under grant DMS-1929284 while one of the authors were in residence at the Institute for Computational and Experimental Research in Mathematics (ICERM) in Providence, RI, during the Theory, Methods, and Applications of Quantitative Phylogenomics semester program. SR was partially supported by the Institute for Foundations of Data Science (IFDS) through NSF grant DMS-2023239 (TRIPODS Phase II). SR was also supported by NSF grant DMS-2308495, as well as a Van Vleck Research Professor Award and a Vilas Distinguished Achievement Professorship.

## A Proof of Eq. (1)

In this section we prove some claims made in Section 2.2.1.

To see why Eq. (1) holds, it suffices to consider the problem of distinguishing species trees with 3 leaves. Let *S* be a 3-leaf species tree topology 12|3, internal branch length *f*, and height *d* > *f*, and let *S*^⋆^ be a star tree of the same height. Suppose one has data consisting of *m* i.i.d. gene trees drawn according to the multispecies coalescent from a 3-leaf species tree 𝒮. Any method capable of estimating a 3-leaf species tree from this data must be able to distinguish between the following two hypotheses:

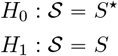

This problem becomes harder as *f* →0 since the two trees become more similar, so as *f* shrinks we expect the data requirement to grow. The key issue is the possible detection of an internal branch, information about which can be ascertained only through the observed coalescence times of lineages 1 and 2 among the *m* gene trees. Under *H*_0_, these two lineages may coalesce only after time *d*, whereas under *H*_1_, they may also coalesce during the time interval (*d* − *f, d*) (see Fig. 6). By elementary coalescent calculations (see below), one can show that if the true species tree parameter is *S*, then the probability of 𝒜^*c*^ is *e*^−*mf*^, and that for any statistical test for distinguishing the two hypotheses, it holds that

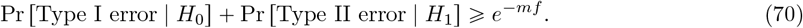

If *m* = *O*(*f* ^−1^), the right-hand side tends to 1 as *f* →0. Hence, the number of genes *m* required to estimate the species tree with high probability must grow at least as fast as *f* ^−1^. The same conclusion holds for trees with *n* 3 leaves since estimating such a tree with minimal internal branch length *f* requires that one be able to resolve all of its 3-leaf subtrees, and the above argument implies that at least one of them—namely, one with internal branch length *f* —cannot be resolved with high probability if *m* = *O*(*f*^−1^). This implies the lower bound in Eq. (1). The takeaway is that no tree with minimal branch length *f* can be reliably estimated if *m* ≪*f*^−1^, since in that case the right-hand side of Eq. (70) is close to 1, meaning that any decision rule which uses the *m* samples as data to distinguish the two hypotheses will be about as good as flipping a coin.

An upper bound matching the lower bound in Eq. (1) can be achieved. Continuing the 3-leaf example, consider a test which rejects *H*_0_ if and only if 𝒜 is observed (i.e., when the minimum observed coalescent time between lineages 1 and 2 among all genes trees is less than *d*). In that case, the combined error rate is

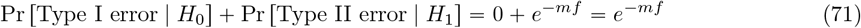

(see below for details). This error can be controlled by increasing *m*; namely, for any *ϵ* >0, the right-hand side is at most ϵ so long as 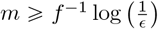. In other words, to distinguish *H*_0_ and *H*_1_ to any pre-set level of power and specificity using this test, it is sufficient that the number of samples satisfies

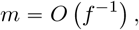

and this upper bound matches the lower bound of Eq. (1). In a similar vein, the GLASS algorithm, a distance-based method which also uses minimum coalescent times from the *m* gene trees, but which can be used to estimate trees with any number of leaves, has been shown to return the correct tree topology with high probability provided that *m* = *O*(*f*^−1^) [15]. Eq. (1) implies that this is the best achievable bound in the limit as *f* → 0, up to multiplication by a constant.

**Proof of Eq. (70) (lower bound):** Let *T* ^(1)^, …, *T* ^(*m*)^ be *i*.*i*.*d*. gene trees drawn according to the multi-species coalescent with species tree parameter 𝒮. Let 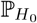 and 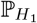 denote the distributions of *T* ^(1)^ under hypotheses *H*_0_ and *H*_1_ respectively. Let *F* = *F*(*T* ^(1)^, …, *T* ^(*m*)^) be any test statistic for distinguishing *H*_0_ and *H*_1_, and let ℛ be the rejection region of *F*. We are interested in controlling the sum of the type-I and and type-II error probabilities

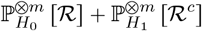

in the setting where *m* → 8 and *f* →0.

As noted in the text, the key issue for distinguishing *H*_0_ and *H*_1_ relates to the detection of the internal branch of *S*, information about which can be ascertained only through the observed coalescence times of lineages 1 and 2. Under 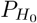, these two lineages may coalesce only after time *d*, whereas under 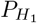, they may also coalesce during the interval (*d* − *f, d*). Let 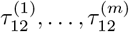 denote the coalescence time of lineages 1 and 2 in the respective gene trees (*T* ^(1)^, …, *T* ^(*m*)^), and define

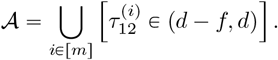

In words, 𝒜 is the event that at least one gene tree does not exhibit ILS on the internal branch. Correctly distinguishing *H*_0_ and *H*_1_ depends crucially on the probability of the event 𝒜; this imposes a data requirement on the size of *m*, which can be made precise as follows:

Under *H*_1_, it holds that 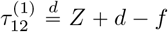, where *Z* is a rate 1 exponential random variable. Therefore, since *T* ^p1q^, …, (*T* ^(1)^, …, *T* ^(*m*)^)are *i*.*d*.*d*.,

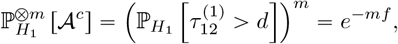

so that

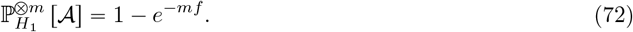

Therefore, in order for any test to reliably distinguish between *H*_0_ and *H*_1_, it is necessary that *m* be sufficiently large that 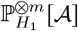 close to 1. By the law of total probability,

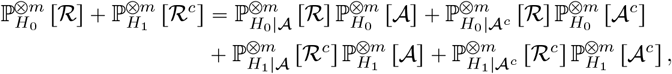

where 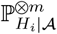 and 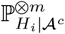 denote the distributions of 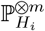 conditional on 𝒜 and 𝒜^*c*^ respectively (*i* ∈ {0,1}).

Since 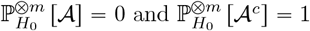, it holds that 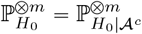. Using these observations along with Eq. (72), it follows that

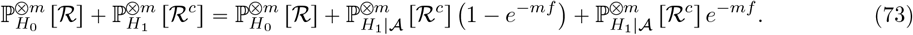

Let 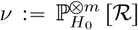. By the Markov property of the multispecies coalescent process, the distribution 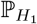 conditional on 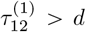 equals the unconditional distribution of 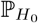. Consequently, 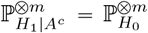, and hence 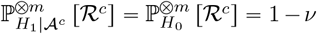. Making these substitutions into the right-hand side of Eq. (73) gives

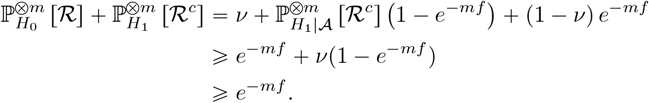

This proves Eq. (70)

**Proof of Eq. (71) (upper bound):** If at least one gene tree is observed with lineages 1 and 2 coalescing prior to time *d*, then one should to reject *H*_0_ since this event has probability 0 under 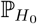. So a natural test to consider is the test with rejection region ℛ = 𝒜. In that case, we can use Eq. (72) to compute the combined error rate:

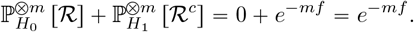

## B Proof of Lemma 1

*Proof of Lemma 1*. Let 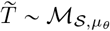 and let 𝒯 ∼ℳ_𝒮,1_. We need to show that 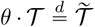. We will do this by showing that for every *x* > 0 with *g*(*x*) >0, the conditional density of 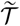, given *θ* = *x*, equals the conditional density of *θ* · 𝒯, given *θ = x*; it will then follow that their expectations with respect to *μ*_*θ*_ must also therefore be equal, i.e., they have the same probability density function.

We begin by considering a single population 𝒳 of 𝒮. Suppose 𝒳 has start and end times *τ*_𝒳_ and *τ*_*P*_ respectively, such that *n* lineages enter 𝒳 at time *τ*𝒳 and *m* lineages exit at time *τ*_*P*_, where 1 ⩽*n* ⩽*m*. Denote by 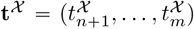 the random vector of *m* − *n* coalescent waiting times in population 𝒳, and let 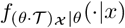 denote the conditional density of the part of the tree *θ* · 𝒯 in population 𝒳, given *θ* = *x*. Then for any **s** = (*s*_*n*+1_, …, *s*_*m*_) ⩾**0** with *s*_*n*+1_ + *s*_*n*+2_ + …+*s*_*m*_ ⩽ *τ*_*P*_ − *τ*_*𝒳*_,

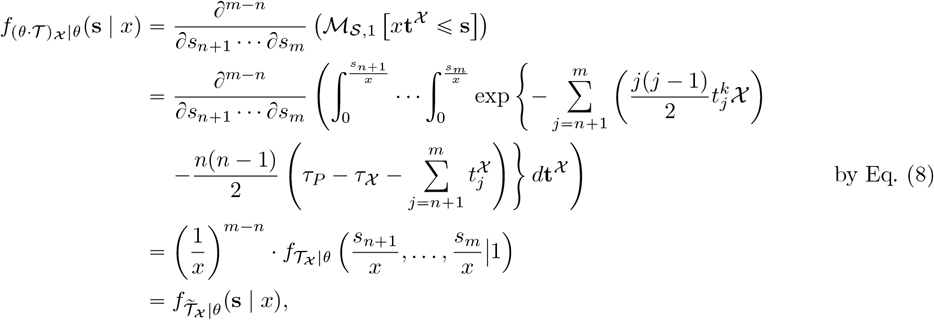

where the second-to-last equality holds by the fundamental theorem of calculus and the chain rule, and the last equality holds by inspection of Eq. (8).

Since 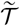 and *θ* · 𝒯 have the same conditional density in each population 𝒳 of 𝒮, it follows by factorization (viz. Eq. (9)) that for every *x* > 0, the conditional densities of 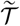 and *θ* · 𝒯, given *θ* = *x*, are equal. Therefore for every gene tree having coalescent waiting times 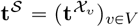,

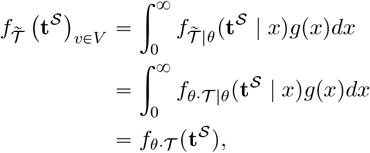

and this implies 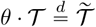.

## C Information-theoretic lemmas

The next two lemmas summarize two important properties of total variation distance which we will utilize in this paper.

### Lemma 13

(Mappings reduce total variation distance). *Let X and Y be random variables taking values in (S*, 𝒮), *let h be a measurable map from (S*, 𝒮) *to (S*^1^, 𝒮^1^), *and let μ*_*W*_ *denote the distribution of the random variable W. Then the following inequality holds:*

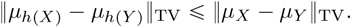

The proof of this lemma follows directly from the definition Eq. (11); see Lemma 4.1.19 in [39].

### Lemma 14

(Subadditivity for product measures). *Consider two probability measures* 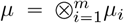 *and* 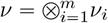 *which are product measures on the same measurable space. Then*

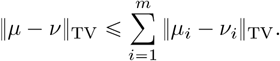

This result is equivalent to Lemma B.8(i) in [40] (see also [41]), and can be easily proved using a coupling argument together with a union bound.

Next, we prove Lemma 2.

*Proof of Lemma 2*. The proof relies crucially on the following *tensorization property* (see, e.g., [36]):

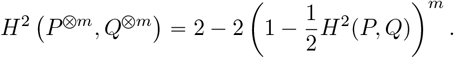

Together with Eq. (13), this implies

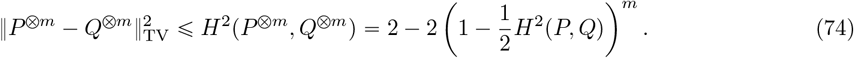

Moreover, by the mean value Theorem 2 − 2(1 −*x*)^*m*^ 2*m*(1 − *ξ*)^*m*−1^*x* for some *ξ* ∈ (0, *x*). Since *m* ⩾ 1, it follows that for all *x* ∈ [0,1],

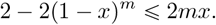

Since 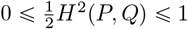 trivially by Eq. (12), we make take 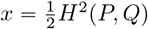 in the above equation, giving

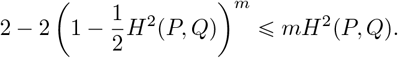

Combining this inequality with Eq. (74) implies the statement of the lemma.

## D Proof of Lemma 4

*Proof of Lemma 4*. By Taylor’s theorem, there exists a number *ξ* between *r* and 1 such that

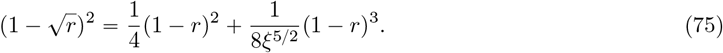

There are two cases, depending on whether *r* > 1 or *r* < 1:

- Case 1. If *r* > 1 then the second term on the right-hand side of Eq. (75) is negative, so

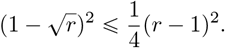 Therefore, by the assumption that *r* ⩽1 + *C*_*E*_*f*,

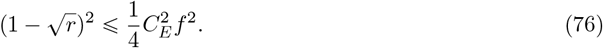
- Case 2. If *r* < 1 then by Eq. (75), along with the assumption that *r* ⩾ 1 − *f*,

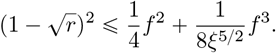

Since *ξ* − *r* and *r* − 1 − *f* − 1/2, it follows that

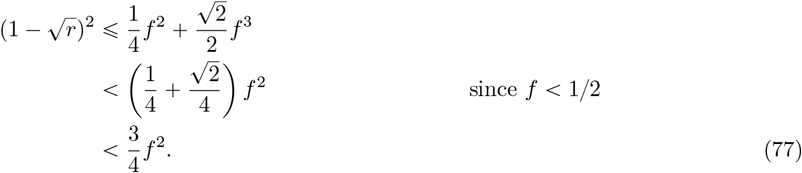

The statement of the lemma follows immediately by Eqs. (76) and (77).

## E Proof that *h* is measurable

### Lemma 15.

*The function h defined in Section 4.4 is measurable*.

*Proof*. Let 𝒜_1_× …×𝒜_*m*_ ∈ 𝒢 ^⊗*m*^. To show that *h* is measurable, it will suffice to show that *h*^−1^ [𝒜_1_ × … × 𝒜_*m*_] ∈ ℱ_*m*_. For each *i* ∈[*m*], let

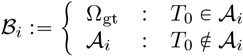

Then

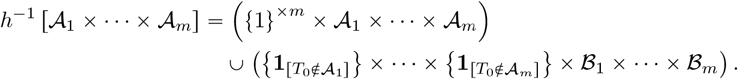

This set is in ℱ_*m*_ since it is a union of two sets which are themselves in ℱ_*m*_.

## Footnotes

1 The *topological distance* between two vertices of a tree is the number of edges on the path connecting them.

